# From Hodgkin-Huxley to Pretrained Neural Inference AI

**DOI:** 10.64898/2026.07.13.738120

**Authors:** Yimu Zhang, Dongqi Han, Zhenning Lv, Fuqing Ren, Yansen Wang, Yang Yang, Dongsheng Li, Yu Gu

## Abstract

High-density probes record from thousands of neurons simultaneously, yet resolving single-neuron identity remains an ill-posed inverse problem. While detailed simulations precisely characterize the biophysical forward process, their utility for interpreting brain signal remains unclear. Here we show that biophysical simulations of population neuronal electrical signals serve as an effective bridge between theory and experiment. By pre-training artificial neural networks exclusively on large-scale synthetic data, we demonstrate robust zero-shot generalization across diverse brain regions, experimental paradigms and species, enabling the accurate inference of single-unit activities and cell-type properties without exposure to real data. Furthermore, uncovering a substantial population of functionally competent but weakly active neurons systematically obscured by conventional heuristics, our framework resolves a long-standing discrepancy regarding ocular dominance in mouse primary visual cortex. These findings establish biophysical simulations as a reference standard, bridging the gap between theoretical understanding and experimental observation through data-driven inference.

## INTRODUCTION

Electrical signals are the language of brain. For decades, scientists have sought to interpret this activity through experiments and theories. From Galvani to Hodgkin-Huxley, these efforts have yielded precise equations for neuronal firing^1–4^ which were further assembled into biophysically and morphologically detailed reconstructions^5,6^. These models capture the mechanisms of individual neurons with biophysical precision, which ostensibly suggests that we have understood the forward problem about how individual neurons “speak a word” of the language. However, when neuron populations “speak” together and are observed by neuroscientists via electrodes, the resulting mixture of signals creates a mathematically ill-posed inverse problem. Despite the precision of these forward models, they have not been effectively utilized to address this inverse challenge. Consequently, a central question remains unresolved: can neural simulations ever be useful for interpreting real brain signals?

Nowadays, modern high-density probes can record millisecond-resolved voltage traces from thousands of neurons simultaneously^7–10^. While we can now obtain much more neural information, this complexity makes the inverse problem more challenging, as the underlying ground truth, including precise spike times, cell identities, and biophysical parameters, remains inaccessible. As a result, every stage of the existing analytical pipeline, from spike detection to cell-type inference, relies on heuristic assumptions. Spike detection typically depends on amplitude thresholds or template matching procedures that are sensitive to noise and parameter choices^11–14^. Subsequent clustering operates on reduced feature representations, often assuming that neurons form stable and separable clusters in a static feature space. In practice, electrode drifting, overlapping waveforms from synchronous firing, and non-stationary biophysical properties routinely violate these assumptions. Even state-of-the-art automated pipelines frequently require manual curation, introducing subjective bias and limiting cross-laboratory standardization^15,16^. Downstream analyses, including cell-type classification, therefore must be built on hand-crafted features derived from these sorted units, with little opportunity for direct experimental validation^17–20^. In this sense, extracellular recordings are not merely noisy; their underlying structure is fundamentally unobservable.

Here, inspired by the transformative success of simulation-to-real (Sim2Real) transfer in controlling complex physical dynamics^21–23^, we address this challenge by leveraging biophysically detailed simulations to pretrain a fully observable artificial intelligence (AI) framework for neural signal inference. Our simulations integrate heterogeneous neuronal morphologies, biophysically grounded electrical properties, and realistic recording physics, yielding a white-box environment in which all latent variables are accessible by construction. This framework enables the generation of domain-randomized synthetic datasets that span the variability encountered in real experiments, allowing us to investigate whether a model trained exclusively on simulated recordings generalize to real biological data.

We show that this simulation-pretrained framework achieves zero-shot neural inference across multiple species and experimental paradigms, accurately performing spike sorting and cell-type prediction without exposure to biological recordings or dataset-specific fine-tuning. Furthermore, we apply this framework to address a long-standing discrepancy regarding ocular dominance in the mouse primary visual cortex. By enabling unbiased, robust inference to single-unit activity, our analysis reveals an intermediate population structure that reconciles conflicting reports from classical electrophysiology and calcium imaging, while also uncovering a substantial population of functionally competent, weakly active neurons that are systematically under-represented by conventional analytical pipelines. Together, these results demonstrate that biophysical simulations serve as a mechanistic and quantitatively foundation for autonomous, generalizable, and self-validating neural inference. Our study advocates a transition from heuristics-driven analysis toward sim-to-real, data-driven neural inference.

## RESULTS

### A simulation-driven, fully observable framework for extracellular electrophysiology

In extracellular electrophysiology, the recorded extracellular potential serves as the primary window to monitor neural activity. However, this aggregate voltage signal is not the ultimate quantity of interest for circuit-level analysis. Instead, essential information resides at the level of individual neurons, including precise spike trains and distinct cell identities^24^. These intrinsic single-neuron variables are not directly observable in the raw recordings, but are mixed into composite extracellular signals and must be recovered through analytical decomposition.

This setting gives rise to two fundamental challenges. First, experimental recordings generally lack ground-truth labels^16,25,26^. Without access to the intracellular membrane potentials of every recorded cell, the true spike sorting solution remains unknown. Second, inferring individual neuronal sources constitutes a challenging inverse problem. Although the biophysical forward process linking neuronal activity to measured voltages is well-defined^27^, mapping noisy and overlapping extracellular signals back to their sources remains mathematically ill-posed.

In contrast to current popular methods which rely on human-designed rules^15^, we propose a simulation-driven deep learning framework, in which large-scale biophysical simulations are used to define ground-truth available training objectives to guide model learning and inference (Fig. 1). The foundation of this framework is the quantitative, comprehensive white-box reconstruction of the electrophysiological environment. We generated large-scale high-density extracellular signals by combining multi-compartment neuronal models^5,6,28,29^ featuring realistic electrical properties and morphologies (Fig. 1A, B). Intracellular spiking activity was first simulated using the NEURON simulator^30,31^, driving the neurons with injected noise currents to ensure realistic responses. Extracellular potentials were then derived using volume conduction theory by summing transmembrane currents across compartments^32^, yielding virtual recordings corresponding to a randomly positioned Neuropixels probe^7,33^. Crucially, this simulation explicitly resolves the geometric relationship between neurons and probes. By computing the Euclidean distance *D*_*i, j*_ between each electrode contact *i* and neuron *j* (Fig. 1B), we defined the Occupancy Intensity *O*_*i, j*_, a variable that quantifies the spatial contribution of neuron *j* to every channel *i*. This variable provides a uniquely defined spatial ground truth which is essential for source separation but fundamentally inaccessible in experimental data. The simulation outputs include complete ground truth annotations, such as spike times, spatial occupancy maps, and cell-identity metadata. These labels enable the inverse problem of extracellular signal analysis to be formulated as a data-driven pretraining task.

**Figure 1.**
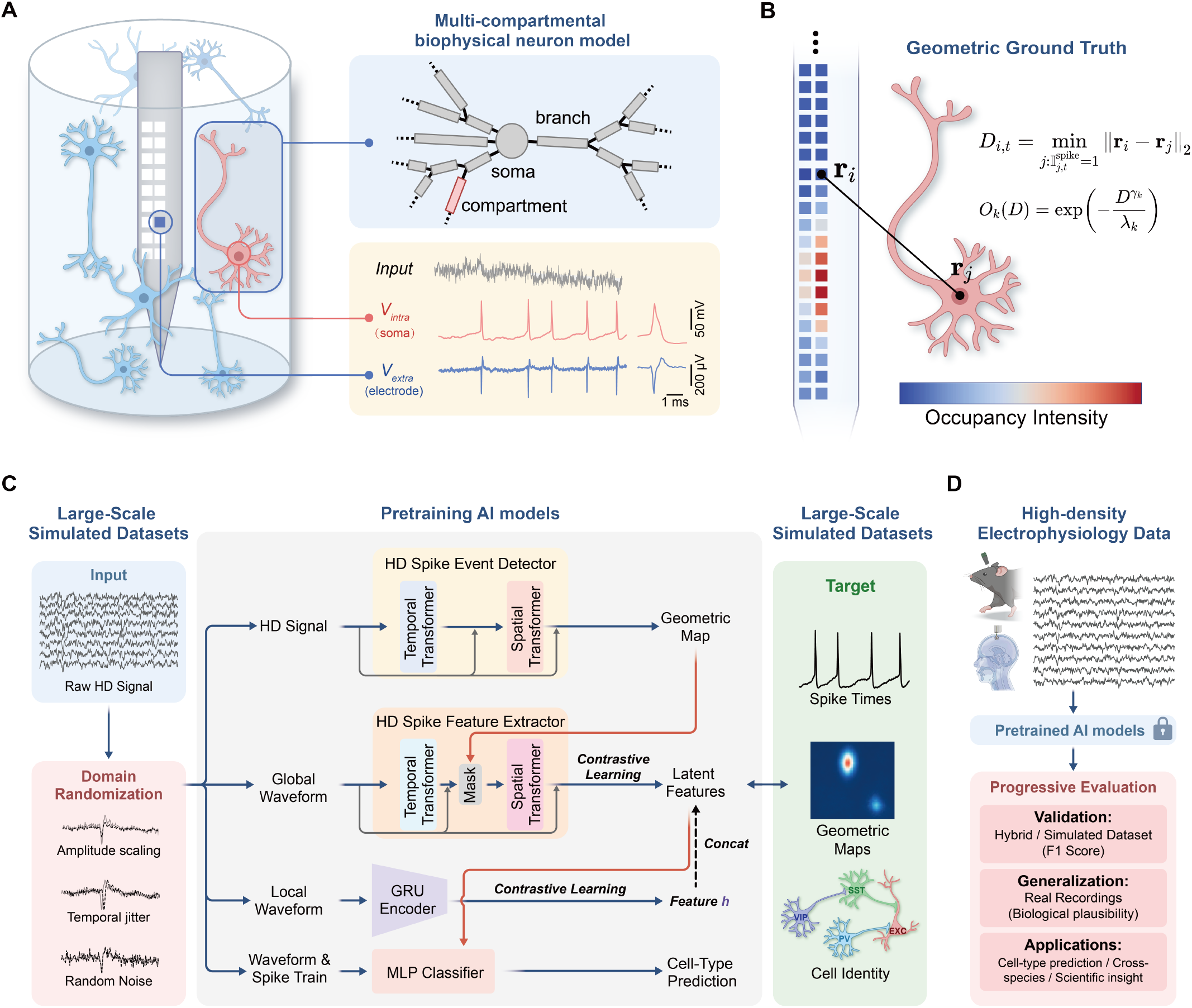
Biophysical simulations provide a fully observable ground truth to train AI models for neural inference. **(A)** Schematic of the simulation environment construction. Multi-compartmental neuron models with realistic morphologies are positioned within a virtual tissue volume alongside a high-density probe. By injecting noise current into the neuron soma, coupled intracellular (*V*_*intra*_) and extracellular (*V*_*extra*_) potentials are generated. **(B)** Definition of geometric ground truth. The Occupancy Intensity (*O*_*k*_(*D*)) is computed based on the Euclidean distance (*D*_*i,t*_) between neuron and electrode contacts. **(C)** Framework architecture and pretraining workflow. Large-scale simulated datasets undergo domain randomization to ensure robust generalization across diverse experimental conditions. These data are utilized to pretrain task-specific deep neural networks: a spatiotemporal Transformer for spike detection supervised by auxiliary spatial maps, dual feature extractors (HD and Local) optimized via contrastive triplet loss to capture both probe geometry and fine waveform morphology, and a multilayer perceptron for cell-type classification that integrates these representations for biological inference. **(D)** The pretrained models undergo a progressive evaluation, progressing from quantitative validation on synthetic datasets to zero-shot generalization on real-world recordings across species, ultimately enabling downstream scientific applications such as cell-type prediction and revealing biological scientific insights.

Based on these electrophysiology simulations, we pretrained a set of AI models for spike event detection, feature extraction, and cell-type inference (Fig. 1C, model architectures detailed in Methods). High-density electrophysiological signals are processed to estimate spike timing and spatial occupancy, while spike waveforms from local and global channel groups are encoded into complementary feature representations. A contrastive objective encourages consistency between features derived from different spatial views. At inference time, these pretrained modules are assembled into a cohesive pipeline applied directly to real recordings: the detection model first localizes spike events from continuous signals, which are subsequently mapped by feature extractors to latent spaces for clustering, and finally assigned biological identities by the classifier (Fig. 1D). To systematically validate the efficacy and generalization capability of this simulation-based framework, the following results present a progressive evaluation ranging from quantitative verification on hybrid datasets to zero-shot applications on diverse experimental recordings across different brain regions, behavioral tasks, and species.

### Quantitative verification on hybrid electrophysiology recordings

Before applying the framework to real experimental recordings, we perform a quantitative evaluation using hybrid neuropixels dataset to assess generalization beyond our simulation domain. This dataset combine real Neuropixels recordings from the International Brain Laboratory (IBL)^34^ with synthetically injected spike events derived from well-isolated units identified by Kilosort4. Waveforms from these units are inserted as additional spike events at spatial positions that are vertically offset from their original detection sites^15^.

We applied the pretrained framework to the hybrid dataset without any fine-tuning. Based on the model-predicted spatial occupancy, spike timing and spatial location were inferred from continuous extracellular recordings (Fig. 2A). Quantitative evaluation revealed strong performance in spike identification and clustering when assessed on the injected ground-truth spike events (Fig. 2B), indicating that the learned representations support accurate unit discrimination. When evaluating the full spike sorting pipeline, including detection, identification and clustering, robust performance was retained despite increased background complexity (Fig. 2C), with 73% of units in the hybrid KS042 recording achieving an F1 score greater than 0.5.

**Figure 2.**
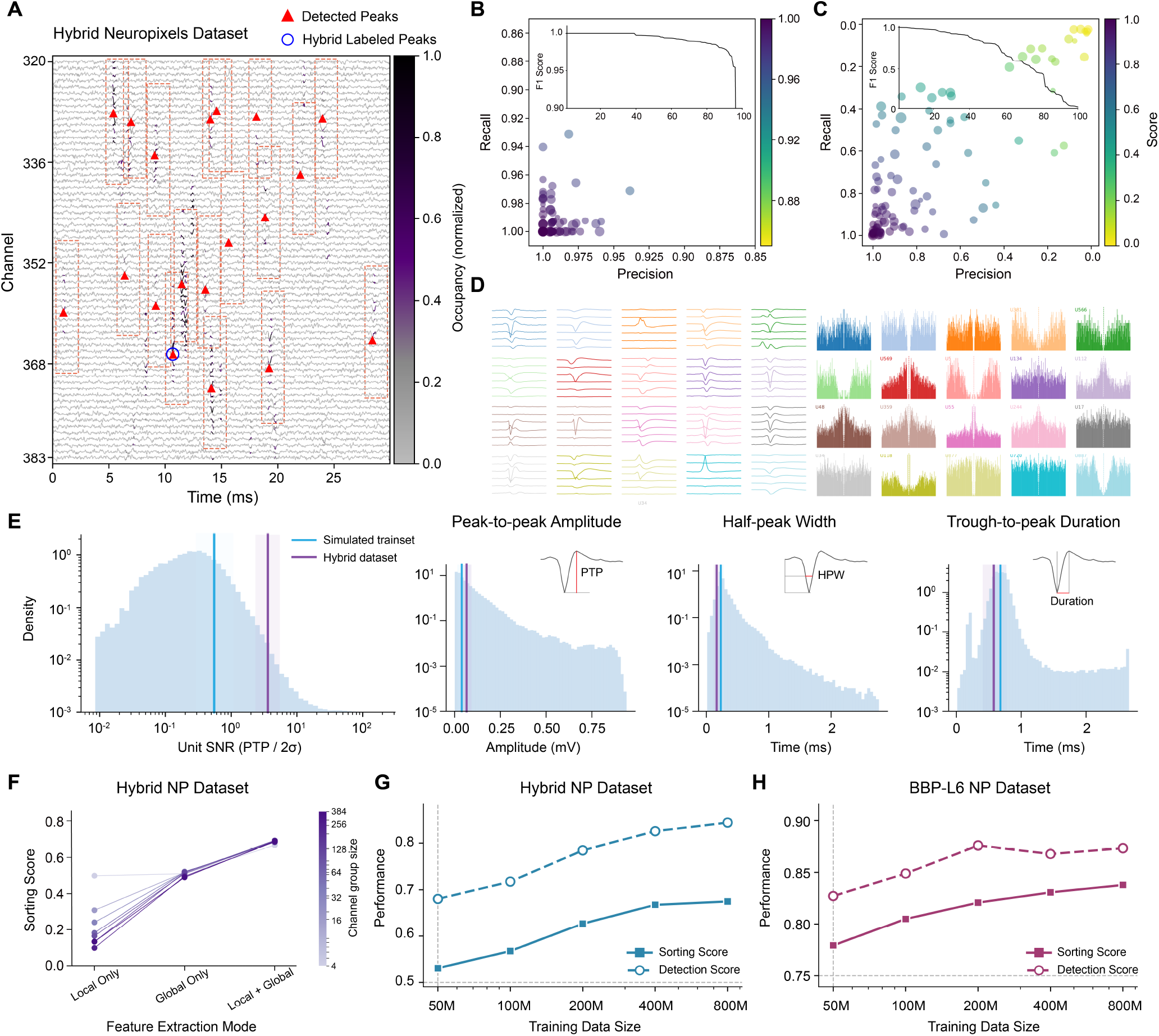
Simulation-pretrained models achieve robust spike sorting on hybrid datasets and demonstrate data scaling laws. **(A)** Visualization of detection inference on the Hybrid Neuropixels recording. The model detects spikes (red triangles) within background activity based on spatio-temporal occupancy, matching injected hybrid labelled peaks (blue circles). **(B)** Evaluation of spike identification and clustering on injected ground-truth spike events. Main plot: Scatter plot of Recall vs Precision for ground-truth units. Inset plot: F1-Score curve across ground-truth units. **(C)** Evaluation of full spike sorting pipeline (spike identification and clustering on detected spikes). Main plot: Scatter plot of Recall vs Precision for ground-truth units. Inset plot: F1-Score curve across ground-truth units. **(D)** Representative waveforms (left) and autocorrelograms (right) of example sorted units. **(E)** Histograms comparing signal statistics (Unit SNR, Peak-to-peak Amplitude, Half-peak Width, and Duration) between the simulated training set (blue) and the hybrid dataset (purple). **(F)** Comparison of sorting scores for different feature extraction modes (Local only, Global only, Local + Global) across varying clustering channel group sizes. **(G-H)** Scaling laws of data size. Sorting and detection scores plotted against the size of the simulated training dataset (50M to 800M samples) for Hybrid NP dataset (**G**) and BBP-L6 dataset (**H**).

Representative examples of identified units showed neuronal-like waveform morphologies and autocorrelograms with clear refractory period (Fig. 2D), illustrating the quality of the putative neurons. Analysis of signal statistics further showed that the simulated training data span a broad range of signal-to-noise ratios and wave-form characteristics, with hybrid recordings largely falling within this distribution (Fig. 2E), enabling the observed simulation-to-hybrid generalization.

One may wonder the underlying reasons of the generalizability of our framework. Here we argue that the contributions of architectural components and data scale are the keys. We first evaluated the performance of independent local (4-channel) and global (384-channel) extractor against a combined architecture across varying channel group sizes used for clustering (Fig. 2F). The results demonstrate that the joint *Local + Global* configuration consistently outperforms single-mode baselines. Furthermore, we investigated the scaling behavior of the framework with respect to training data volume. By varying the size of the simulated dataset from 50 million to 800 million samples, we observed a clear monotonic improvement in both detection and sorting performance on Hybrid NP and BBP-L6 datasets (Fig. 2G, H, Figure S1). This scaling law indicates that the model effectively capitalizes on the vast diversity of the simulation, suggesting that the performance ceiling has not yet been reached and can be further elevated by scaling the synthetic corpus.

### Zero-shot generalization on real electrophysiology recordings

After quantitative validation using synthetic datasets, we evaluated the framework on fully experimental electrophysiology recordings. Since the ground truth of individual spikes is unavailable, performance was assessed using functional and behavioral signatures in two real datasets: one from a visual coding task^10,35^, and the other from a decision-making task^36,37^.

We first applied the framework to the Allen Visual Coding Neuropixels dataset, which comprises standardized visual stimuli recorded from the mouse primary visual cortex (Fig. 3A). Based on the sorting results inferred in a zero-shot manner, we analysed receptive field structure (Figure S2A), stimulus-evoked responses (Figure S2B), and orientation selectivity (Figure S2C). Representative example units exhibited clear stimulus-locked firing patterns and well-defined tuning properties across stimulus conditions (Fig. 3B-D), consistent with canonical visual cortical responses^38,39^.

**Figure 3.**
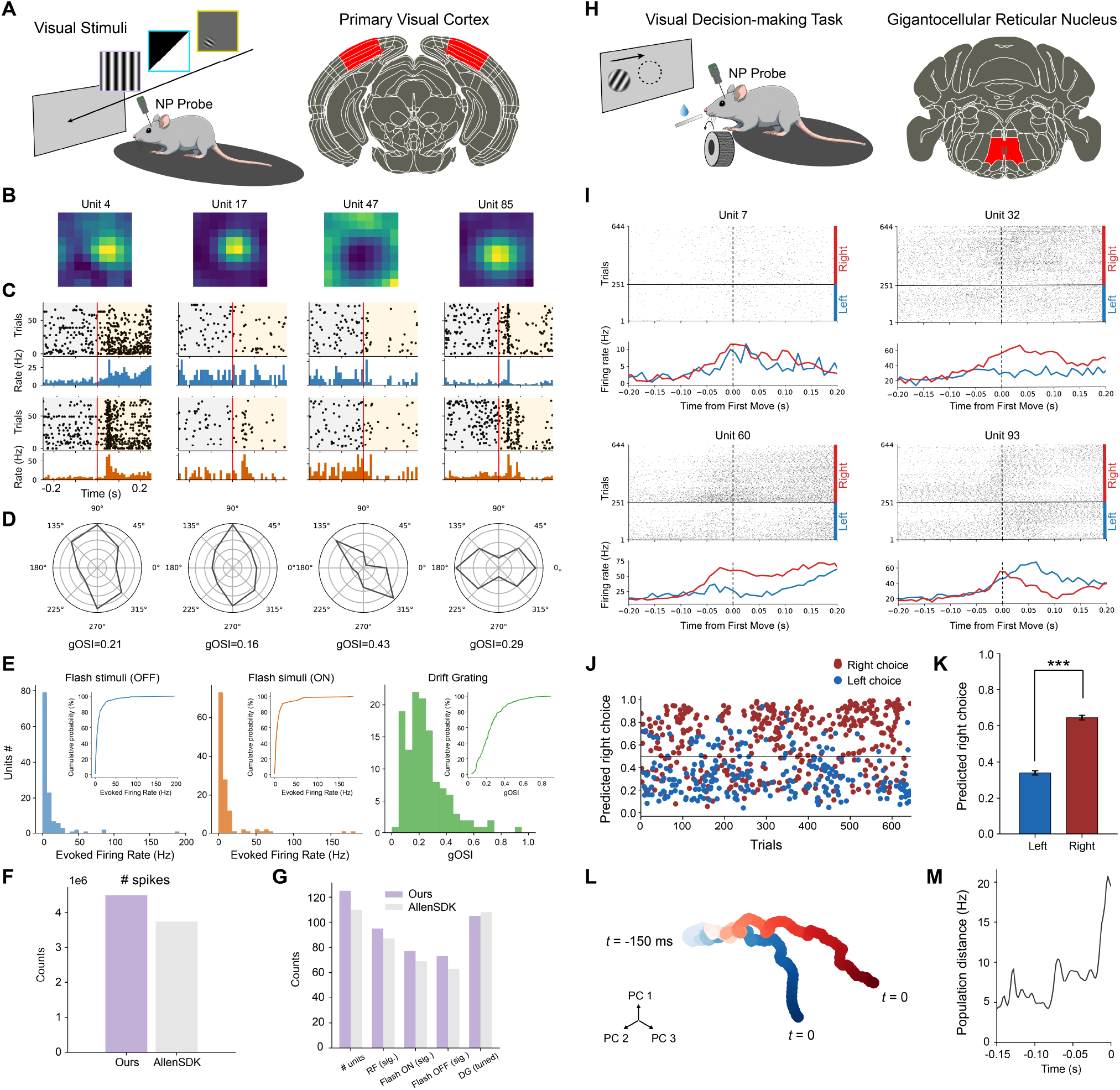
Zero-shot generalization to real electrophysiology recordings recovers biological plausible sensory and behavioral representations. **(A)** Schematic of the experimental setup for the Allen Visual Coding dataset (V1 recordings). **(B)** Visual receptive fields of four representative units inferred by our framework. **(C)** Raster plots and PSTHs aligned to Flash OFF/ON stimuli for the example units shown in **B. (D)** Orientation tuning curves for the example units showing orientation selectivity. **(E-G)** Comparison of population statistics between the simulation-pretrained framework and the AllenSDK. **(E)** Main plots show histograms of evoked response magnitudes and global Orientation Selectivity Index (gOSI); Insets show the corresponding Empirical Cumulative Distribution Functions. **(F)** Total spike counts. **(G)** Counts of total units, counts of units with significant receptive field, counts of units with significant ON response, counts of units with significant OFF response, and counts of units with gOSI greater than 0.1. **(H)** Schematic of the experimental setup for the IBL decision-making dataset. **(I)** Raster plots and PSTHs of example units aligned to the decision period (Left vs. Right choice). **(J-K)** behavioral decoding analysis. **(J)** Predicted right choice probability over trials. **(K)** Average decoding accuracy for left and right choices. **(L-M)** Neural population dynamics. **(L)** PCA trajectories coloured by time relative to choice. **(M)** Euclidean distance between left- and right-choice population trajectories over time.

To assess reliability at the population level, we compared these results with the official analyses provided by the Allen Brain Observatory through the AllenSDK, which are based on Kilosort2 outputs followed by expert curation. Across stimulus conditions, our framework recovered comparable numbers of visually responsive units and produced highly consistent response profiles, including evoked response magnitudes and orientation selectivity distributions (Fig. 3E-G). This close agreement indicates that the neural activities obtained without dataset-specific tuning support functional characterizations aligned with established reference analyses.

We next examined zero-shot generalization in a behavioral context using the IBL decision-making dataset. Recordings were obtained from the gigantocellular reticular nucleus (GRN) while mice performed a visually guided wheel-turning task, choosing left or right in response to visual cues (Fig. 3H). The framework identified single units modulated by the animal’s choice during the decision period (Fig. 3I), indicating task-related encoding at the single-unit level.

At the population level, neural activity derived from the identified units predicted the animal’s upcoming choice (Fig. 3J, K). Moreover, population state trajectories corresponding to left and right choices diverged progressively in a low-dimensional representation, with the Euclidean distance between trajectories increasing during the decision epoch (Fig. 3L, M). These population dynamics are consistent with previous reports of choice-related representational separation in decision-making circuits^37^.

Together, these results show that single-unit activity inferred by the simulation-pretrained framework preserves established sensory and behavioral structure in real electrophysiological recordings across brain regions and tasks. This consistency indicates generalization beyond simulated and hybrid domains.

### Simulation data-driven cell-type classification *in vivo*

Beyond resolving the spatiotemporal dynamics of spiking activity, determining the biological identity of recorded neurons is essential for dissecting neural circuit mechanisms. We therefore asked whether our framework can recover biologically meaningful cell-type distinctions from extracellular signals.

To address this question, we extended our framework to cell-type classification as a downstream inference task. Using biophysically detailed neuron models from the Allen Cell Types database^40^, we pretrained a classifier on simulated datasets with labels for transcriptomic cell classes (excitatory, PV, SST, and VIP) and morphological spiny types (spiny versus aspiny). Representative examples of modelled neurons illustrate the diversity of morphologies, extracellular waveforms, and firing dynamics captured by the simulations (Fig. 4A). The simulated dataset spans distinct waveform features and firing-pattern statistics across cell types, including differences in spike shape and inter-spike interval distributions (Fig. 4B, C), indicating that the simulations encode discriminative structure relevant to cell identity.

**Figure 4.**
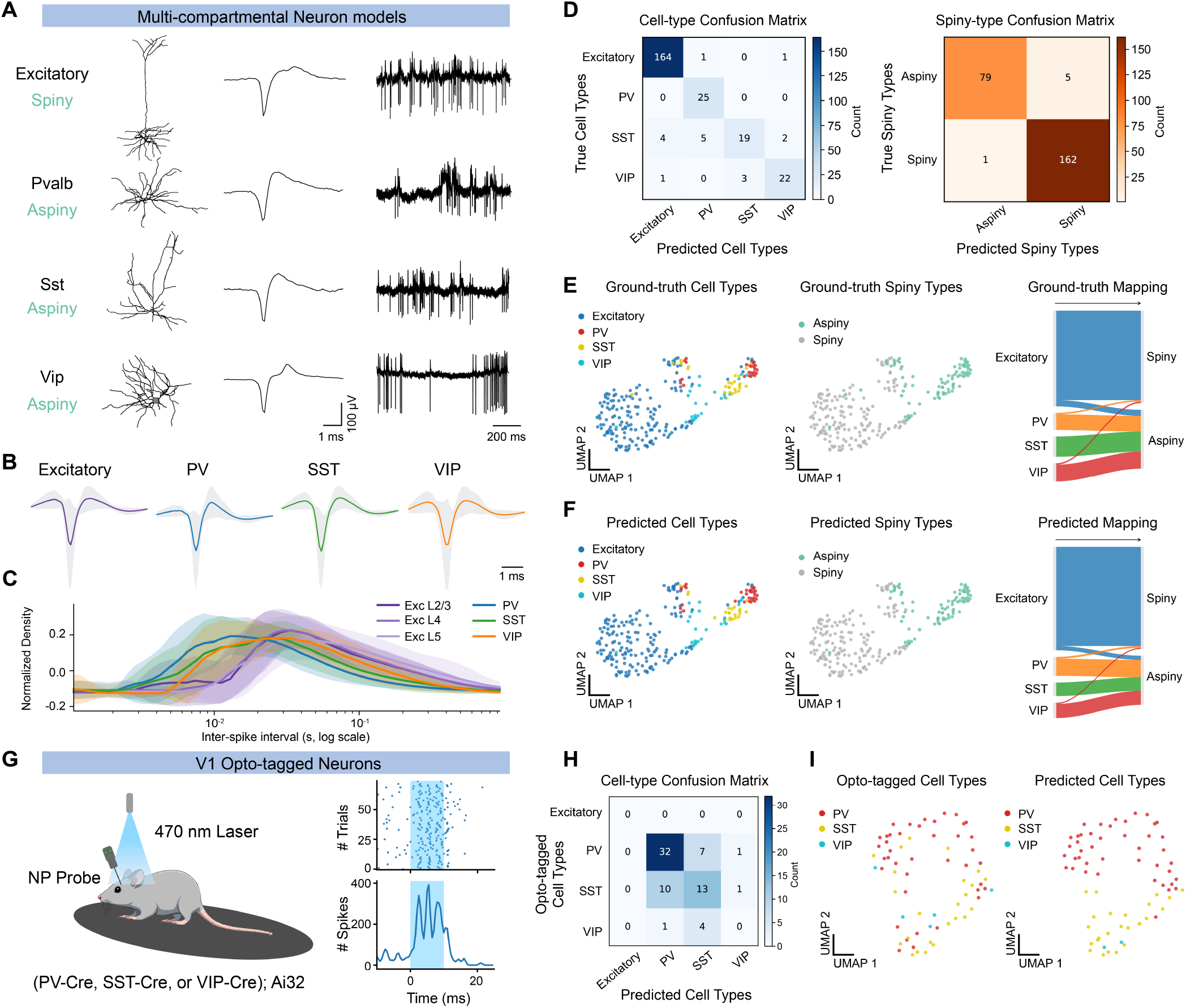
Simulation-based pretraining extracts biologically meaningful cell-type features verifiable *in vivo*. **(A)** Visualization of simulated neuron models for four cell types (Excitatory, Pvalb, Sst, Vip), showing morphology, extracellular waveforms, and raw trace. **(B)** Mean extracellular waveforms (coloured lines) overlaid with standard deviation (grey shading) for distinct cell types (Excitatory, PV, SST, VIP) in the simulated dataset. **(C)** Normalized density distributions of log-transformed inter-spike intervals (ISIs) for the corresponding cell types in the simulated dataset. **(D)** Confusion matrices for cell-type (left) and spiny-type (right) classification on the held-out simulated test set. **(E)** UMAP projection of the latent feature space coloured by true cell-type labels (left), true spiny-type labels (middle), and Sankey diagram mapping true cell types to spiny types (right). **(F)** UMAP projection of the latent feature space coloured by predicted cell-type labels (left), predicted spiny-type labels (middle), and Sankey diagram mapping predicted cell types to spiny types (right). **(G)** Schematic of the optogenetic tagging validation on real data (PV, SST, VIP Cre-lines). **(H)** Confusion matrix for cell-type classification on real opto-tagged neurons. **(I)** UMAP visualization of real opto-tagged neurons (left) and their predicted identities (right).

We first evaluated the classifier on held-out simulated test data. The model achieved high accuracy in predicting both transcriptomic cell types and spiny types (Fig. 4D). Confusion matrices revealed clear separation between excitatory and inhibitory neurons, as well as among inhibitory subtypes. Low-dimensional embeddings of the learned representations exhibited clear organization by true cell type and spiny type, with predicted labels closely aligned with the underlying ground-truth structure (Fig. 4E, F). Sankey diagrams summarizing the correspondence between predicted cell classes and spiny types further illustrate the consistency between functional classification and morphological organization.

We next evaluated whether the simulation-pretrained classifier generalizes beyond simulated data to real extra-cellular recordings. To obtain cell-type ground truth, we analysed optogenetically tagged Neuropixels recordings from mouse primary visual cortex in Allen visual coding dataset^10^, using Cre-line targeting PV, SST, and VIP inhibitory neurons. By selecting neurons that responded reliably to laser stimulation, we constructed a test set of opto-tagged inhibitory cells (Fig. 4G). When applied to these recordings in a zero-shot manner, the classifier correctly identified the opto-tagged neurons as inhibitory, with particularly high accuracy for PV cells (Fig. 4H, I).

Collectively, these results demonstrate that representations learned from simulations support cell-type inference that generalizes to real experimental data. A classifier trained purely on simulated recordings can recover biologically meaningful distinctions in both simulated and real neuronal populations, validating simulation-based pretraining as an effective strategy for resolving neuronal identities in real electrophysiology recordings.

### Cross-species generalization to human extracellular recordings

The scarcity of human extracellular recordings presents a long-standing, dual challenge. On one hand, the comprehensive interpretation of neuronal activity in these rare datasets is scientifically critical to maximizing their value. On the other hand, the development and validation of analytical methods for human data are substantially constrained by this very scarcity and the unavailability of definitive ground-truth annotations. Fortunately, the neurons share many common properties between humans and rodents, providing the potential for our framework to generalize from mice and rats to humans^41^.

Given the lack of direct validation metrics, we evaluated performance by assessing the physiological consistency of the inferred spike trains through qualitative analyses of human cortical recordings acquired under anesthesia from an epilepsy patient (Pt03)^42^.

Single-unit spike trains were extracted using our framework, and time-locking analyses were conducted to examine the coupling between neuronal spiking activity and prominent LFP (Local field potential) events. We focused on two temporally structured phenomena present in this dataset: anesthesia-associated burst suppression and epilepsy-related interictal epileptiform discharges (IIDs) (Fig. 5A, B).

**Figure 5.**
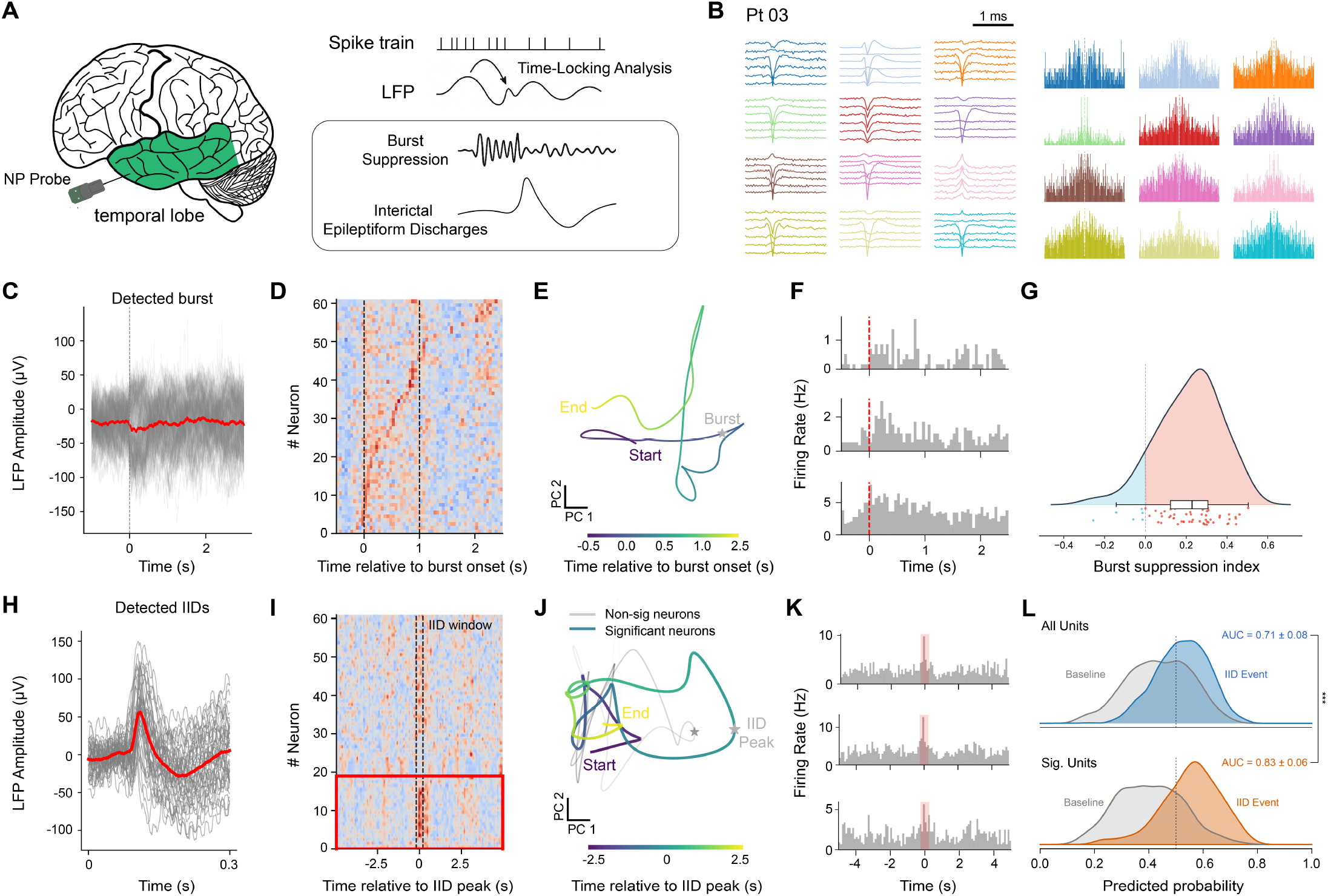
Representations learned from mouse simulations generalize cross-species to human cortical dynamics. **(A)** Schematic of human temporal lobe recordings and the analysed LFP events: Burst Suppression and Interictal Epileptiform Discharges (IIDs). **(B)** Waveforms and autocorrelograms of example units sorted from human data (Pt 03). **(C)** Traces of LFP waveforms of burst events. **(D)** Heatmap of single-unit firing rates aligned to burst onset. **(E)** PCA trajectory of population activity during burst suppression. **(F)** PSTHs of three example neurons aligned to burst onset. **(G)** Distribution of the burst suppression index for the recorded population. **(H)** Traces of LFP waveforms of detected IIDs. **(I)** Heatmap of single-unit firing rates aligned to IID peaks. The red box indicates the population of neurons exhibiting significant response during IID window. **(J)** PCA trajectories of IID-significant (teal) and non-significant (grey) neuronal populations. **(K)** PSTHs of three example neurons aligned to IID peaks. **(L)** ROC curves quantifying the detection of IID events based on population spiking activity. Curves represent the mean performance of decoders trained on all units (blue) and significant units (orange) across n=50 independent bootstrap iterations. Shaded regions indicate standard deviation. The significant units demonstrated significantly higher decoding accuracy compared to the full population (p<0.0001, paired two-sided *t*-test).

During burst suppression, large-amplitude LFP bursts were interspersed with periods of marked quiescence (Fig. 5C). Aligning neuronal firing to burst onset revealed distinct modulation patterns: while some neurons exhibited surge-like firing during the burst phase, others showed suppression (Fig. 5D, F). At the population level, neuronal activity evolved along a coherent trajectory over time (Fig. 5E). We quantified this effect using a burst suppression index, which revealed substantial functional heterogeneity across the recorded population (Fig. 5G).

We next examined neuronal responses to interictal epileptiform discharges (IIDs). Detected IIDs exhibited stereotyped LFP waveforms with sharp peaks (Fig. 5H). Time-locking spike trains to these peaks identified a subset of neurons with temporally precise firing increases centered on the event (Fig. 5I, K). Population dynamics evolved along a coherent trajectory during the IID window, driven primarily by these responsive units, whereas non-responsive units showed minimal excursion (Fig. 5J). To quantify the informational content of the spiking activity, we assessed the ability of single-neuron spike trains to discriminate LFP time windows containing IIDs using a Random Forest decoder. ROC analysis demonstrated that the IID-responsive neurons achieved significantly higher detection performance compared to the full population (paired *t*-test, p<0.0001; Fig. 5L), confirming robust spike-LFP coupling during epileptiform activity.

To complement these functional analyses with biological identification, we applied our cell-type classifier trained solely on mouse simulations to the human recordings to infer transcriptomic cell types and spiny morphologies. The structural organization of these predicted populations is visualized in the model’s latent embedding space (Figure S3A-C). The inferred cell-type assignments exhibited expected morphological biases, with excitatory units preferentially associated with spiny morphologies and inhibitory subtypes (PV, SST, VIP) biased toward aspiny profiles (Figure S3B, F). We further observed that the degree of IID modulation varied across these predicted cell types (Figure S3E), raising the possibility that distinct inhibitory subtypes may be differentially recruited during pathological discharges.

Moving forward from the previous sections, which demonstrates that generalizability from simulation to reality, cross experiment tasks, and cross brain-regions, the results of humans recordings further suggest that our framework, trained exclusively on mouse simulations has the potential to generalize to human cortical recordings. The observation that inferred units exhibit dynamics consistent with prominent physiological events and preserve biologically meaningful cell-type signatures supports the feasibility of using simulation-based pretraining for cross-species analysis.

### Revealing biological insights into ocular dominance in mouse V1b

Having demonstrated the framework’s robustness and zero-shot generalization capabilities in complex human cortical recordings, we next sought to apply its unbiased inference power to resolve specific biological questions. While generalization across species validates the model’s biophysical foundation, its scientific utility is ultimately defined by its ability to uncover neuronal populations that are systematically missed by conventional heuristic methods. To test this, we focused on a specific analytical challenge where method-dependent sampling bias has historically obscured the ground truth. A long-standing discrepancy exists regarding ocular dominance in the binocular zone of mouse primary visual cortex^43^. Classical electrophysiological studies generally describe a predominance of binocular neurons with ocular dominance indices clustered around middle values^44–47^, whereas several two-photon calcium imaging studies have reported a substantial fraction of monocular neurons^48–51^(Fig. 6C). The origin of this divergence remains unclear.

**Figure 6.**
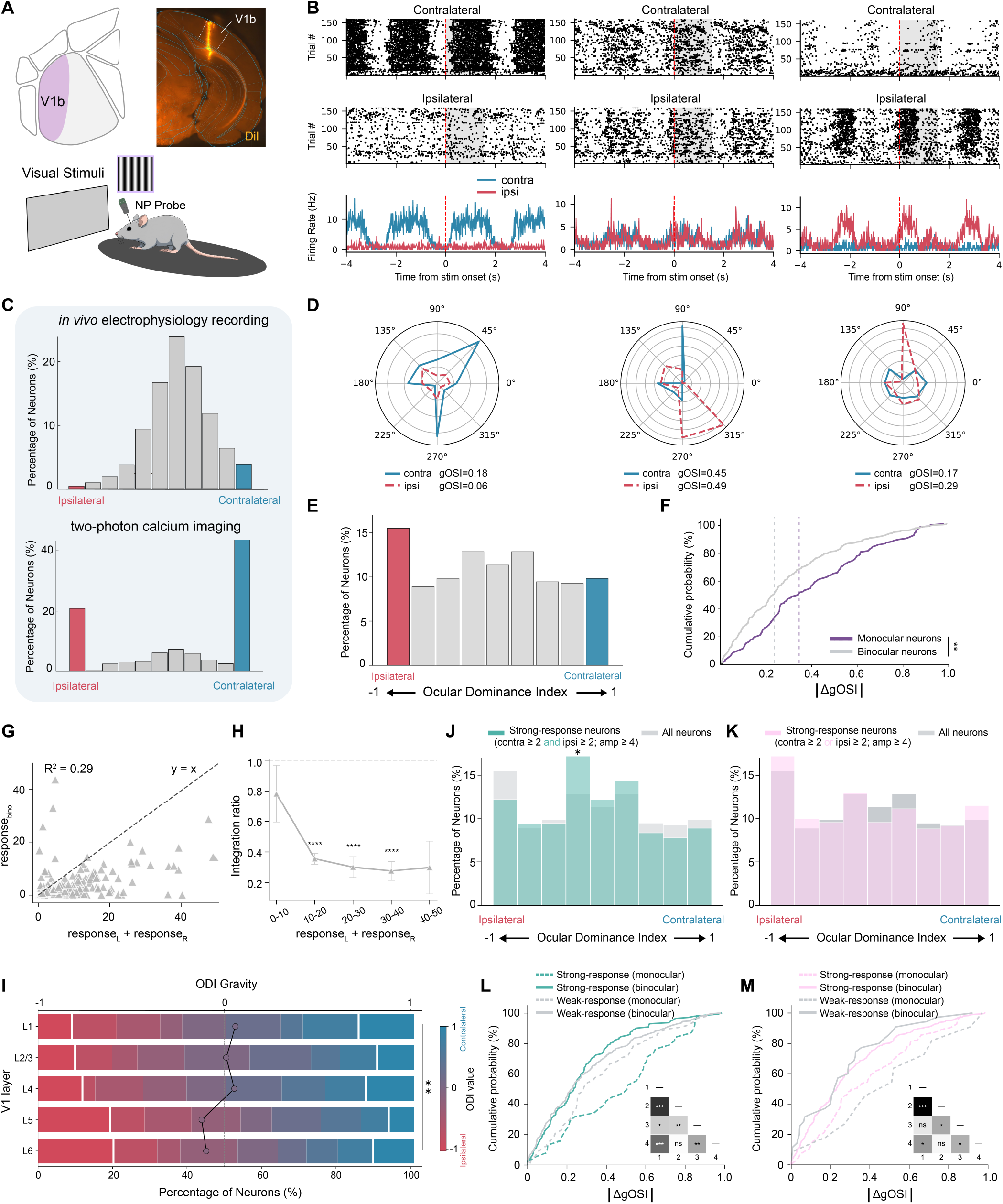
The simulation-pretrained framework enables unbiased inclusion of weakly active neurons, reconciling conflicting ocular dominance profiles in mouse V1. Figure 6. **(A)** Schematic of the recording setup in the binocular zone of mouse V1 with DiI track verification (N = 7 mice). **(B)** Raster plots and PSTHs of three example neurons (Contralateral, Ipsilateral, Binocular) under monocular and binocular stimulation. **(C)** Schematic comparison of ocular dominance (OD) distributions reported by electrophysiology vs. two-photon imaging. **(D)** Orientation tuning curves for the example neurons. **(E)** Histogram of the Ocular Dominance Index (ODI) derived from the framework (n = 528 units). **(F)** Cumulative probability plots of the difference in global orientation selectivity (|ΔgOSI|) for monocular and binocular neurons. **(G)** Scatter plot of binocular response versus the linear sum of monocular responses. Dashed line represents unity. (H) Integration ratio plotted against response magnitude bins. **(I)** ODI values plotted across V1 cortical layers (Spearman rank correlation, p=0.008). **(J)** ODI distribution for “strong-response neurons” selected using “AND” criteria (responsive to both eyes). Statistical comparison performed against the remaining population (p = 0.039, Fisher’s exact test). Waveform and autocorrelogram examples of the weak-response neurons are shown in Figure S4. **(K)** ODI distribution using “OR” criteria (responsive to either eye). **(L)** Cumulative probability of |ΔgOSI| for strong-response (coloured) and weak-response (grey) neurons under the “AND” criterion. Solid and dashed lines denote binocular and monocular neurons, respectively. The inset displays the *p*-value matrix for pairwise comparisons; indices 1–4 correspond to the legend entries from top to bottom (*p* values, Mann-Whitney U test). **(M)** Cumulative probability of |ΔgOSI| under the “OR” criterion. The inset shows the corresponding *p*-value matrix for pairwise comparisons.

Here, our framework unveils the potential causes of this discrepancy. We applied our simulation-pretrained framework on acute Neuropixels recordings from the binocular zone (Fig. 6A). By enabling spike detection and unit inference without parameter tuning or extensive post hoc curation, our framework mitigates the sampling biases and allows for a more objective analysis. We first assessed the physiological validity of the inferred single-unit activity. Representative units extracted by the framework exhibited distinct spiking patterns (Fig. 6B) and selective orientation tuning (Fig. 6D) under monocular and binocular stimulation.

Interestingly, the population-level distribution derived from our analysis did not conform exclusively to either historical extreme. Instead, it occupied an intermediate regime (Fig. 6E). Physiologically, this observed distribution is supported by functional properties: binocular neurons exhibited significantly tighter interocular global orientation selectivity matching (|ΔgOSI|) than monocular neurons (Fig. 6F) and displayed sublinear integration (Fig. 6G, H), consistent with previous reports^52^. Furthermore, we observed a significant layer-dependent shift in ocular dominance (Fig. 6I). This trend is consistent with mouse V1 retinotopy^53^, as vertical probe insertions into arcuate brain surface likely sample more lateral, ipsilateral-dominant coordinates at greater depths.

We hypothesized that the long-standing conflict stems from sampling bias toward high-amplitude units in electrophysiological analyses. To test this, we simulated the selection criteria of classical electrophysiological studies by restricting analysis to neurons with strong responses to *both* eyes (the “AND” criterion) and high waveform amplitudes. This constraint artificially shifted the distribution to reproduce the classical dominance of binocular neurons (Fig. 6J). In contrast, we broadened the criteria to include neurons responsive to *either* eye (the “OR” criterion). This approach mimics the logic of calcium imaging analysis pipeline. Under this inclusive condition, the bias disappeared, and the resulting distribution recovered the monocular populations at the extremes, matching the full population (Fig. 6K).

Crucially, we verified that these recovered weak-response units represent a distinct but functionally competent population. Within the binocular category, response strength did not predict the precision of binocular integration; weak-response neurons exhibited orientation tuning consistency identical to that of strong-response neurons (Fig. 6L, M, insets). Moreover, the functional distinction between weak- and strong-response monocular neurons, which appeared significant under restrictive criteria, disappeared under the inclusive “OR” condition (Fig. 6M).

These findings suggest that weak-response monocular neurons are functionally competent units with tuning properties comparable to strong-response neurons. While traditional extracellular analyses often miss these units due to amplitude-based thresholds and low signal-to-noise ratios, our results demonstrate that they constitute a genuine physiological population. Resolving this long-standing debate highlights the critical utility of our simulation-pretrained framework. By enabling the unbiased recovery of neurons obscured by conventional limitations, it reconciles the divergent findings of electrophysiology and imaging, providing a unified view of visual cortical organization that encompasses the full spectrum of neuronal responsiveness.

## DISCUSSION

Deciphering the activity of individual neurons from extracellular recordings is a mathematically ill-posed inverse problem. The absence of ground-truth labels *in vivo* has led most existing approaches to rely on heuristic rules^11,14,15,54,55^. Here, we address this fundamental limitation by presenting a simulation-driven AI framework that learns the inverse mapping of the biophysical signal generation process. Leveraging a fully observable white-box environment, we pretrain deep neural networks to recover latent neuronal activity from spatiotemporal voltage patterns. Our results demonstrate that models trained exclusively on biophysically grounded simulations achieve zero-shot generalization to real recordings, robustly spanning diverse species and experimental conditions.

Zero-shot generalization in this framework emerges from the combination of conserved biophysical laws and domain randomization. This strategy aligns with a broader paradigm in scientific machine learning, where models trained on physics-based simulations have successfully solved inverse problems in fields ranging from nuclear fusion plasma control to robotic manipulation^21,56,57^. Just as agents trained in virtual physics engines can generalize to physical hardware, our work demonstrates that biophysical simulations can serve as a robust foundation for sim-to-real transfer in electrophysiology. By randomizing neuronal morphology, spatial organization, and noise structure across a wide parameter range, we constrain the network to learn invariant causal relationships, ensuring that real experimental recordings fall within the subspace covered by the simulated data.

The universality of these biophysical principles enables extension from rodent models to human cortical recordings, where labelled data and direct validation are typically unavailable. In Fig. 5, we show that models trained on rodent simulations are possible to decode spike timing and cellular identity in human brain. This result reflects shared principles of bioelectric signal generation and potentially supports the use of simulation-pretrained models as standardized tools for clinical neuroscience. The fully observable nature of the simulation framework also allows access to latent variables that cannot be measured *in vivo*, enabling future extensions to circuit-level inference.

Our application of this framework helps reconcile the disparate ocular dominance profiles reported by electrophysiology and calcium imaging. By recovering weakly responsive neurons that are typically underrepresented in conventional extracellular analyses, we reveal an intermediate population structure consistent with both observations. Functional analyses in Fig. 6 show that tuning properties are preserved across response strengths, indicating that weakly responsive neurons are not qualitatively distinct from strongly responding ones. These results suggest that method-dependent sampling bias, rather than differences in neuronal tuning, accounts for a substantial component of the reported discrepancy.

Scaling this simulation-driven approach points to extensive future work. Future iterations must expand the simulation library to encompass a broader spectrum of neuronal morphologies and biophysical properties. Specifically, incorporating distinct electrophysiological signatures, such as the complex spikes of cerebellar Purkinje cells, will be essential for robust generalization across diverse brain regions.

By combining explicit biophysical modeling with advanced model design and simulation-based pretraining, this framework links microscopic biophysical processes to circuit-level observations and supports unbiased analysis across recording conditions and species. Ultimately, by anchoring AI in biophysical simulations, this framework defines an elegant pathway for mechanistic inference in data-scarce neuroscience problems, with implications for translational neuroscience and brain-computer interfaces where ground truth is fundamentally inaccessible.

## METHODS

### Animals

C57BL/6J wild-type mice were used in this study (n = 7). All animals were male, aged 8-10 weeks, and weighed 20-25 g. Mice were obtained at approximately 7 weeks of age from Shanghai Lingchang Biotechnology Co., Ltd. The sample size (number of animals) was predetermined to include at least three mice with recordings from either the left or right primary visual cortex (V1), in order to ensure unbiased statistical analysis of the expected neuronal population. Prior to surgery, mice were group-housed in a specific pathogen-free (SPF) animal facility under a 12 h light/dark cycle, and experiments were performed during the light phase of the cycle. The temperature ranged from 19-23 °C and the humidity was 55% ± 10%. Mice had free access to food and water. The mice were single housed in cages (37×15×13 cm) after surgery. All experimental procedures were conducted in full compliance with institutional guidelines and were approved by the Animal Ethics Committees of Fudan University and ShanghaiTech University.

### *in vivo* Electrophysiology recordings

#### Surgical procedures

Upon arrival, mice were allowed to acclimate to the housing environment for 3 days. On day 4, mice were anesthetized with isoflurane (3% for induction and 1.5–2% for maintenance) and secured in a stereotaxic frame. The scalp was shaved, incised, and the periosteum was removed to expose the skull. The skull surface was cleaned using 0.9% saline and 3% hydrogen peroxide. After skull exposure, bregma and lambda were identified and marked, and the head position was adjusted to align the anteroposterior and mediolateral axes. Using bregma as a reference, the target location of the primary visual cortex (AP: −3.7 mm, ML: ± 2.75 mm) was marked on the skull. In addition, a small burr hole was drilled above the cerebellum, into which a skull screw was implanted to serve as the ground and reference electrode for Neuropixels recordings. Subsequently, a custom-made stainless steel headplate was implanted and secured to the skull using dental cement to enable head fixation. To protect the skull without obscuring the V1 target markings, a thin layer of dental cement was applied over the skull surface. On day 9, a 1-mm-diameter craniotomy was performed at the marked V1 location using a high-speed dental drill. After craniotomy, the opening was sealed with biocompatible silicone elastomer (Kwik-Cast™, World Precision Instruments) and UV-curable adhesive. On day 10, mice were handled for approximately 30 minutes to facilitate habituation prior to the onset of Neuropixels recordings.

#### Visual stimuli

Visual stimuli were generated using custom-written scripts in MATLAB and presented via the Bpod State Machine (Sanworks; https://github.com/sanworks/Bpod_Gen2). Stimuli were displayed on a Dell e1715s monitor (resolution 1280 × 1024, screen width 34 cm, refresh rate 60 Hz) positioned 15 cm in front of the mouse. During the experiments, visual stimuli were presented sequentially under binocular and monocular viewing conditions. For monocular recordings, the non-recorded eye was occluded with black opaque tape applied using silicone oil, and the eye was subsequently cleaned with sterile saline following the recording session. The visual stimulus consisted of full-field drifting sinusoidal gratings with spatial frequency of 0.08 cpd, temporal frequency of 2 Hz and contrast of 100%. Gratings were presented in eight different directions of 0^°^, 45^°^, 90^°^, 135^°^, 180^°^, 225^°^, 270^°^, 315^°^, and stimulus directions were presented in a pseudorandomized order. Each stimulus was presented for 1.5 s, followed by a 1.0 s inter-stimulus interval consisting of a uniform black screen (blank condition). Each orientation was repeated 20 times.

#### Neuropixels recording

We performed acute, high-density *in vivo* electrophysiological recordings using Neuropixels 1.0 probes (IMEC, Belgium) to record from mouse primary visual cortex (V1). Neural signals were acquired using a PXI-based data acquisition system (National Instruments) and recorded with SpikeGLX software (https://github.com/billkarsh/SpikeGLX). The amplifier gain was set to 500× for action potential (AP) channels and 250× for local field potential (LFP) channels. Signals were referenced and the AP band was high-pass filtered at 300 Hz.

Prior to each recording, the UV-curable adhesive and biocompatible sealant covering the probe were carefully removed. Probes were inserted using a motorized micromanipulator (Multi-Probe Manipulator, New Scale Technologies) at a speed of 400 *µ*m/min, and the probe tip was positioned at a depth of 1500 *µ*m, ensuring that the recording sites spanned the depth range of V1. To ensure proper grounding of the recording system, the ground wire of the Neuropixels probe was electrically connected to a skull screw via a lead wire. After probe insertion, the probe was allowed to settle in the brain for approximately 5 minutes before the onset of data acquisition.

Each mouse was recorded once under each stimulus condition, including binocular stimulation, left-eye-only stimulation (with the right eye occluded), and right-eye-only stimulation (with the left eye occluded). During recordings, the cortical surface was kept moist using sterile saline applied with a 5 mL syringe. For post hoc verification of electrode placement and reconstruction of probe trajectories, each probe was coated with the fluorescent lipophilic dye DiI (C1036, Beyotime Biotechnology), which allowed visualization of the electrode track and identification of recording sites.

#### Histology and image registration

After completion of all experiments, mice were deeply anesthetized with isoflurane until loss of reflexes was confirmed. Once a stable level of anesthesia was achieved, animals were transcardially perfused with 10 mL of 1× phosphate-buffered saline (PBS), followed by 10 mL of 4% paraformaldehyde (PFA) delivered at a flow rate of approximately 4-5 mL/min. After perfusion, the brains were then carefully extracted and post-fixed in 4% PFA for 12 hours.

Following fixation, brains were sectioned into 80-*µ*m-thick coronal slices using a vibrating microtome (VT1200S, Leica Microsystems) and mounted onto glass slides. Images were subsequently acquired using a fluorescence stereomicroscope (MVX10, Olympus) to verify the location and depth of the Neuropixels probe tracks labelled with fluorescent dye. For visualization and registration of the brain section images, we used the HERB software platform (https://github.com/Whitlock-Group/HERBS). Image registration was performed by aligning the acquired brain sections to the “Allen Mouse Brain Atlas”.

### Electrophysiology simulation

#### Neuron models

Multi-compartmental neuron models were obtained from two publicly available, biophysically detailed databases: the Neocortical Microcircuit Collaboration of the Blue Brain Project^5,58^ and Allen Brain Cell Types^6,40^.

The Blue Brain Project (BBP) models comprise 206 individual neuron reconstructions from the rat somatosensory cortex, spanning cortical layers L1-L6 across 30 distinct cell classes. Each model incorporates up to 13 active ion channel types together with an explicit model of intracellular *Ca*^2+^ dynamics. Axon initial segments (AIS), somata, basal dendrites, and apical dendrites were represented as distinct morphological regions; interneurons contained a single dendritic compartment. Each morphological region was assigned an independent set of ion channels. Model parameters were constrained using extensive in vitro electrophysiological recordings and systematic morpho-electrical classification, yielding multiple morpho-electrical variants that reproduce characteristic firing patterns across cell types^5^.

Allen Brain neuron models (Allen models) were derived from mouse primary visual cortex L1-L6. The models contain 17 cell classes, represented by 112 unique individual neuron models. These models cover diverse excitatory and inhibitory cell classes and were parameterized using experimentally measured intrinsic electrophysiological properties and anatomical features, as described in the data-driven cortical modeling framework^6^.

#### Intracellular simulation

Intracellular electrophysiological activity was simulated using the NEURON package^30,31^. Pink noise, scaled to the rheobase (the minimum current needed to elicit an action potential in a neuron), was injected into each neuron’s soma to evoke stochastic action potentials, simulating biologically realistic background noise. The temporal resolution was set to 0.1 ms, and simulations spanned a total duration of 600 seconds. Neurons were initialized with a resting membrane potential of −70 mV and maintained at a physiological temperature of 34°C. Synaptic mechanisms and biophysical properties were dynamically compiled and loaded to ensure compatibility with the NEURON simulation environment.

For each neuron, a multi-compartmental model^5,6,28,29^ was used to calculate transmembrane currents^32,59–61^. The neuron was divided into compartments, each described as an equivalent electrical circuit. The dynamics of the membrane potential in each compartment *n* were governed by Kirchhoff’s current law^62^, where the sum of all currents entering or leaving a node must equal zero. Considering that extracellular potential changed much slower than ion channel dynamics, we assumed it constant. The temporal evolution of the membrane potential *V*_*m*_ in compartment *m* was given by,

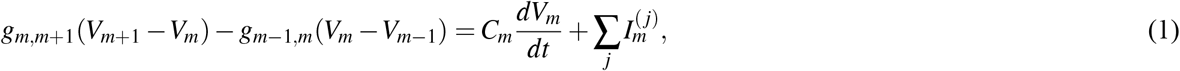

where *g*_*m,m*+1_ and *g*_*m*−1,*m*_ represented the conductances between neighboring compartments, *C*_*m*_ was the membrane capacitance, and 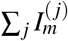 accounted for ionic currents and any externally applied currents.

These transmembrane currents served as the source for extracellular potential modeling in the next simulation stage.

#### Extracellular potential

Extracellular potentials were computed using volume conductor theory, which relates transmembrane currents to extracellular voltages measured at electrode sites^27,63,64^. In each simulation trial, 50 neurons were randomly sampled from the available model pool and positioned within a three-dimensional volume surrounding the electrode, with spatial bounds of −30 to +30 μm along the x-axis, 0 to +50 μm along the y-axis, and −30 to +30 μm along the z-axis. Neurons of multiple types were included to represent the cellular diversity of cortical microcircuits. A virtual high-density electrode with a Neuropixels 1.0/2.0-like site layout was placed within the same volume to record the resulting extracellular potentials.

The extracellular potential at a given electrode position **r**_*e*_ was computed as the linear superposition of contributions from all neuronal compartments within the simulation domain. Under the line source approximation, each compartment was modelled as a finite line segment carrying a uniformly distributed transmembrane current. The extracellular potential was calculated as

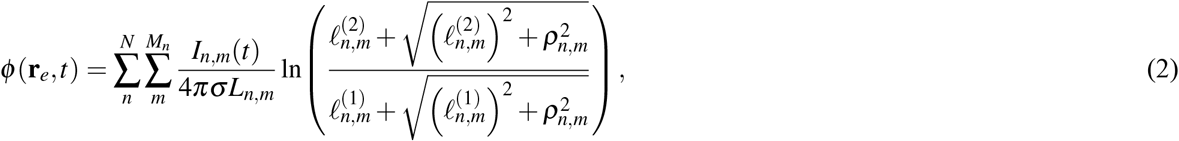

where *I*_*n,m*_(*t*) denotes the transmembrane current of the *m*-th compartment of the *n*-th neuron at time *t*, and *L*_*n,m*_ is the length of that compartment. The geometric quantity *ρ*_*n,m*_ represents the shortest perpendicular distance from the electrode position to the axis of the compartment, while 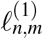 and 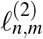 denote the axial distances from the electrode to the proximal and distal endpoints of the compartment, respectively. Here, *N* is the total number of neurons and *M*_*n*_ is the number of compartments of the *n*-th neuron. The extracellular medium was assumed to be infinite, homogeneous, and isotropic with conductivity *σ*. A reference electrode located at infinity was assumed, such that the potential vanishes at infinity.

### Large-scale synthetic datasets

We performed large-scale electrophysiological simulations and generated multiple synthetic datasets for model training and evaluation.

#### Simulated continuous signal dataset

This dataset was constructed from 1,000 trial extracellular recordings simulated using BBP neuron models and Allen models. In each simulation trial, 50 neurons were selected and extracellular potentials were computed at high-density probe sites with a temporal resolution of 0.1 ms. For each simulation output, a fixed-length segment of the continuous extracellular signals was extracted (typically 100,000 time steps for HD-detection model training and 600,000 time steps for evaluation). Extracellular signals were organized as continuous multichannel time series. Spike labels were derived from ground-truth intracellular spike events provided by the simulator. Spike onset times were first identified from the intracellular spike label matrix, and spike peak times were then estimated using a fixed temporal offset. Binary detection labels were constructed by assigning positive labels within a short temporal window surrounding each spike peak. In addition to binary spike labels, a channel-wise distance map was computed to encode the spatial relationship between spike events and electrode sites. For time points containing spikes, the Euclidean distance between each channel and the corresponding neuron soma was calculated, and the minimum distance across active neurons was assigned to each channel at that time point, resulting in a continuous spatiotemporal distance representation aligned with the extracellular signals.

In addition, we generated a layer-restricted dataset (BBP-L6) by constraining neuron sampling to layer 6 models only. During simulation, Blue Brain Project models from layers 1-5 were used exclusively for training, whereas layer 6 models were held out and used only for evaluation.

#### Simulated spike waveform dataset

The spike waveform dataset was constructed from 1,000 trial simulated extra-cellular recordings using intracellular ground-truth spike labels. This dataset was used to train the local extraction model and HD-extraction model. Extracellular signals were band-pass filtered at a sampling rate of 10 kHz. Probe site coordinates and neuron soma positions were used to define spatial relationships between channels and units. A channel-wise occupancy representation was computed from spike events by mapping the minimum Euclidean distance between each probe channel and spiking neuron somata through an exponential kernel. Waveforms were extracted per unit using a fixed 60-sample window across all probe channels.

#### Simulated cell-type dataset

The simulated cell-type dataset was constructed from 200 trial electrophysiological recordings generated using Allen neuron models, which include representative cell types from the mouse primary visual cortex (V1), including excitatory pyramidal neurons and major inhibitory interneuron classes such as parvalbumin-expressing (PV), somatostatin-expressing (SST), and vasoactive intestinal peptide-expressing (VIP) neurons. For each simulated recording, extracellular signals, intracellular spike labels, probe site coordinates, and neuron soma positions were loaded. Cell-type annotations and dendritic spiny labels were obtained from the associated metadata. Neurons with insufficient spiking activity or recording duration were excluded. For each neuron, spike trains as well as waveform and occupancy features were extracted from ground-truth spike events. Firing pattern features were then computed from the spike trains using the logarithmic inter-spike interval distribution.

### Simulation data-driven pretraining framework

#### Overview of the framework

We introduce a simulation-based pretraining framework for high-density extracellular recordings that factorizes neural inference into modular, learnable components. The framework comprises four modular components:

1. a High-Density (HD) spike event detector operating on continuous multichannel signals;
2. an HD spike feature extractor that encodes probe geometry and spatial spike footprints;
3. a local spike feature extractor that captures fine waveform morphology from peak-centered channel subsets;
4. a cell-type classifier that integrates latent representations for biological inference.

Each module is pretrained on large-scale simulated datasets and subsequently composed at inference time to process real recordings without dataset-specific fine-tuning.

#### HD spike event detector

Spike detection is formulated as a dense temporal prediction task on the continuous multichannel voltage signal **V** ∈ ℝ^*T×C*^, where *T* denotes the number of time points and *C* the number of recording channels (*C* = 384 for Neuropixels probes^7,33^). Unlike sparse tetrode recordings, high-density probes exhibit strong spatial correlations. To exploit this, we employ a spatiotemporal Transformer architecture that processes the full multichannel context to output a spike probability score for each time point.

To encourage spatially consistent event localization, the model is trained to predict auxiliary spatial maps (e.g., occupancy intensity) alongside per-timepoint probabilities. The total detection loss is a weighted sum of binary cross-entropy terms across prediction heads:

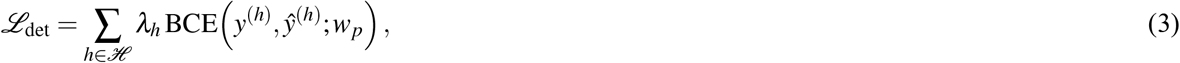

where ℋ represents the set of output heads (temporal probability and auxiliary spatial maps), BCE(·) denotes the class-weighted binary cross-entropy, λ_*h*_ are head-specific balancing weights, and *w*_*p*_ is a positive-class weight handling spike sparsity.

#### HD spike feature extractor

Following detection, candidate events are extracted as fixed-length multichannel snippets *X*_*i*_ ∈ ℝ^*L×C*^ centered on the detected spike time. The HD feature extractor is designed to map these snippets to spatially aware embeddings that explicitly encode Neuropixels probe geometry and the distributed extracellular footprint. In addition to raw voltage traces, the model incorporates spatial inductive biases–such as channel adjacency and occupancy representations–to preserve the relative topological organization of the recording sites. Pretraining relies exclusively on simulated data, allowing direct supervision of spatial structures (e.g., source location) which are inaccessible in real recordings. The model is optimized using a metric learning objective based on triplet loss^65^, encouraging embeddings from the same neuron to cluster tightly while separating distinct units:

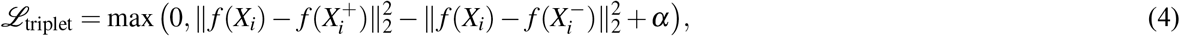

where *f* (·) is the embedding function, 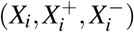 form an anchor-positive-negative triplet, and *α* is the margin and 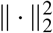 denotes the squared L2 norm. The resulting embeddings are compatible with standard dimensionality reduction methods (e.g., PCA, UMAP) and downstream clustering algorithms.

#### Local spike feature extractor

To complement the global spatial context, a local feature extractor characterizes the fine-scale temporal and morphological properties of spikes. This module operates on a subset of channels selected based on proximity to the event’s dominant channel (peak channel). We adapted the waveform encoder from SimSort^66^, employing a lightweight GRU-based encoder. The model aligns and denoises the local waveforms before projecting them into a compact latent space suitable for distinguishing subtle morphological variations.

#### Cell-type classifier

The classification module integrates the complementary representations from the HD and local extractors. Latent vectors are concatenated and processed by a multilayer perceptron (MLP) to output a probability distribution over predefined neuronal cell types (e.g., Excitatory, PV+, SST+, VIP+).

### Training procedure

#### Optimization and scheduling

All models were implemented in PyTorch and trained on simulated datasets. The HD spike event detector was trained using the AdamW^67^ optimizer. To stabilize training on the highly imbalanced spike detection task, we employed an explicit learning-rate warm-up (4,000 steps) followed by a cosine annealing scheduler. The HD spike feature extractor and cell-type classificier were optimized using AdamW with a step-wise learning rate decay (StepLR). The local spike feature extractor was optimized using Adam^68^ with a StepLR and trained using a triplet margin loss. Anchor-positive-negative triplets were constructed by the dataset during data loading.

#### Domain randomization and augmentation

To ensure generalization from simulation to experiment, we applied comprehensive data augmentations during training. For the detection model and feature extractors, channel-level augmentations included additive Gaussian noise, random amplitude scaling, and temporal jitter, calibrated to match the noise statistics of experimental recordings. These domain randomization strategies prevent overfitting to simulation artifacts and improve zero-shot transfer performance.

### Inference pipeline

During inference, raw signals are first band-pass filtered and spatially whitened. The continuous data is processed in a streaming fashion using overlapping windows with cached context to handle long-duration recordings seamlessly. Each window is passed to the pretrained HD Spike Detector, which outputs multi-head probability maps. Peak detection is performed via a multi-stage consensus process: (i) candidate peaks are identified using 2D non-maximum suppression on Gaussian-smoothed binary maps; (ii) these are cross-validated against the auxiliary occupancy map and raw voltage extrema; and (iii) redundant detections are consolidated within a local spatiotemporal neighborhood using component-based pooling.

For every consolidated event, the system extracts two views: a geometry-aligned snippet for the HD extractor and a peak-centered local snippet for the local extractor. These pretrained models yield global spatial and fine-grained local embeddings, respectively. Subsequently, distinct neuronal units are resolved from these high-dimensional representations using a group-wise clustering strategy (detailed below). Finally, the consolidated unit embeddings are fed into the cell-type classifier to predict neuronal identity. This pipeline supports efficient, parallelized processing and allows for modular integration with downstream analysis steps such as drift tracking and quality control.

#### Group-wise clustering and global merging

To improve clustering accuracy while maintaining computational tractability for large-scale high-density arrays, we adopted a group-wise clustering strategy. The recording probe topology was partitioned into overlapping spatial windows (default spatial window size *W* = 96 channels, with an overlap of *O* = 32 channels). Individual spike events were assigned to windows according to the location of their peak channel.

Within each local window, spike embeddings were projected onto a lower-dimensional manifold using UMAP to enable reliable density estimation. Gaussian Mixture Models (GMMs) were then applied to identify local clusters. By restricting clustering to spatially localized waveforms, this procedure mitigates the noise accumulation and interference effects commonly encountered in global clustering, thereby improving the separation of nearby units.

To assemble coherent global neuronal identities from these local clusters, we implemented a greedy sequential merging algorithm that iterates across adjacent spatial windows. Local clusters were merged into a single global unit if they satisfied two consistency constraints:

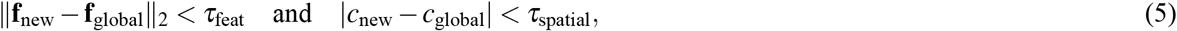

where **f** denotes the cluster centroid in feature space and *c* denotes the mean peak channel. We set the feature similarity threshold to *τ*_feat_ = 0.05 and the spatial proximity threshold to *τ*_spatial_ = 10 channels. This merging strategy ensures that neurons spanning multiple spatial windows are consistently stitched together, while preventing the erroneous fusion of spatially distinct units with similar waveform features.

### Evaluation

#### Hybrid Neuropixels dataset^**15**^

Hybrid electrophysiology recordings released with Kilosort4 were used for quantitative evaluation. These datasets are generated by injecting synthetic spike events into real Neuropixels recordings collected by the International Brain Laboratory (IBL)^34^. The injected events are derived from well-isolated units identified by Kilosort4, whose extracellular waveforms are re-inserted at spatial locations vertically offset from their original detection sites. This procedure preserves the original background activity and noise structure of the recordings while providing spike-level ground-truth annotations for the injected events.

All analyses on hybrid data were performed on the *KS042* recording. Raw broadband signals were processed directly by the inference pipeline without fine-tuning or dataset-specific parameter adjustment. The hybrid dataset was used to evaluate spike detection, identification, and clustering behavior in the presence of injected ground-truth events embedded in real extracellular recordings.

#### Allen visual coding dataset^**10**,**35**^

Neuropixels recordings from the Allen Visual Coding resource were included to evaluate zero-shot inference on sensory-evoked cortical activity. The dataset consists of extracellular recordings obtained from head-fixed mice during standardized visual stimulation paradigms, including drifting and static grating stimuli, with probes targeting primary visual cortex and associated visual areas. One recording (session_id = 732592105; probe_id = 733744649) was analysed. Extracellular signals from the primary visual cortex were processed using the same pretrained inference pipeline as in other datasets, without retraining or parameter tuning. This dataset was used for downstream analyses of visually driven neural responses and population activity derived from spike sorting outputs.

#### IBL decision-making dataset^**36**,**37**^

The IBL decision-making dataset comprises Neuropixels recordings acquired during a visually guided decision-making task. Mice performed a wheel-turning task requiring left or right choices in response to visual cues, while neural activity was recorded simultaneously across multiple brain regions under a unified experimental protocol. One recording (eid: aec5d3cc-4bb2-4349-80a9-0395b76f04e2; pid: 7332e6cf-9847-4aca-b2e3-d864989dd0fb) was analysed. Extracellular signals from the gigantocellular reticular nucleus (GRN) were processed directly by the pretrained inference pipeline without dataset-specific modification. This dataset was used to support downstream analyses of task-related single-unit activity and population-level neural dynamics.

#### Human Neuropixels dataset^**42**^

Publicly available human Neuropixels recordings were analysed to assess cross-species generalization. These recordings were acquired from cortical regions of an epilepsy patient under anesthesia and exhibit increased noise levels, waveform variability, and nonstationary background activity relative to rodent recordings.

Analyses focused on the recording from patient *Pt03*. Raw extracellular signals were processed using the same simulation-pretrained inference pipeline without modification. Due to the absence of spike-level ground truth, this dataset was used exclusively for downstream physiology-driven analyses based on inferred spike trains and population activity.

#### Allen cell classification dataset^**10**^

To evaluate cell-type prediction performance, a labelled dataset was curated from the Allen Visual Coding resource by selecting mouse visual cortex sessions with available optogenetic tagging metadata and excluding wild-type sessions. Putative inhibitory neuron classes were defined using commonly used Cre lines (Pvalb, Sst, and Vip). For each unit, baseline and light-evoked firing rates were computed in fixed temporal windows relative to light onset (baseline: [−100, 0] ms; evoked: [0, 50] ms). Units were identified as opto-tagged using a conservative firing-rate ratio criterion, FR_evoked_*/*FR_baseline_ ≥ 4. Waveform features were extracted from peak-centered, fixed-length multi-channel snippets spanning all probe channels. For each unit, snippets were aligned to the trough on the peak channel, and 60 samples (± 30 samples at 30 kHz) were extracted from all channels. Temporal firing statistics were derived from inter-spike-interval histograms computed on spontaneous activity, using 20 logarithmically spaced bins between 1 and 1,000 ms. All features were z-scored within each session. Labels were assigned according to inhibitory classes defined by Cre-driver lines (PV, SST, and VIP). These data were used exclusively for evaluation, and were not used for training or fine-tuning any model components.

### Definitions of metrics

#### Spike sorting

Spike sorting performance was evaluated by event-level matching between detected spike trains and ground-truth spike trains. Ground-truth spikes were provided as tuples of spike time, unit identity, and a representative channel, whereas detected spikes were represented by spike times with associated cluster labels and detected channels. For a given ground-truth unit *l* consisting of *N*_*l*_ spikes with spike times 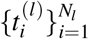 and a detected cluster *k* consisting of *M*_*k*_ spikes with spike times 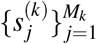, a detected spike was considered matched to a ground-truth spike if their absolute time difference satisfied

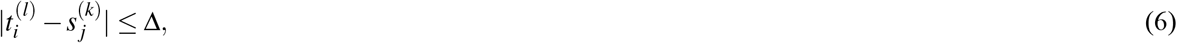

with a matching window of Δ = 1 ms. A one-to-one constraint was enforced at the spike-event level such that each ground-truth spike and each detected spike could be matched at most once. The number of matched events between unit *l* and cluster *k* was denoted as 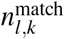, from which true positives, false positives, and missed events were defined

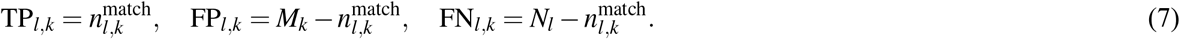

Precision and recall were computed as

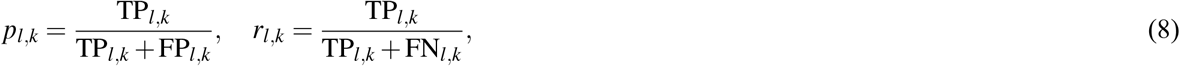

and the F1 score was used as the primary per-unit performance metric:

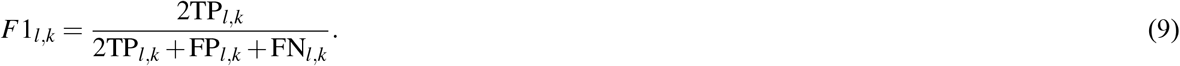

To restrict comparisons to spatially plausible candidates, each detected cluster was assigned a representative channel 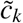 defined as the median detected channel across its spikes. For each ground-truth unit *l* with representative channel *c*_*l*_, only the *K* = 40 clusters with the smallest channel distance 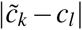 were considered. The best-matching cluster for unit *l* was selected by maximizing the F1 score over this candidate set,

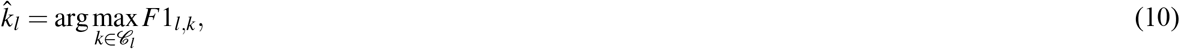

and per-unit precision, recall, and F1 score were reported accordingly. Overall spike sorting performance was summarized as the mean F1 score across all ground-truth units,

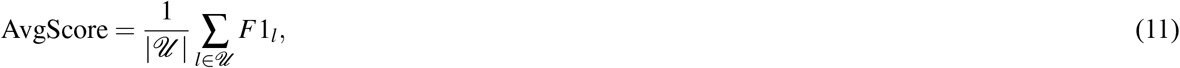

where *U* denotes the set of ground-truth units.

#### Cell-type classification

Cell-type and spiny-type prediction were treated as multiclass classification tasks. Performance was evaluated using classification accuracy,

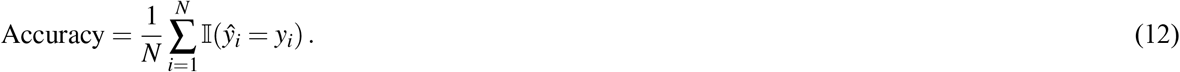

where *N* denotes the total number of samples, and *y*_*i*_ and *ŷ*_*i*_ represent the ground-truth and predicted labels for the *i*-th sample, respectively.

### Data analysis

#### Visual receptive field

Receptive fields (RFs) were mapped using the Gabor stimulus table from the Allen Visual Coding dataset. For each stimulus presentation, spike counts were summed within a response window [*t*_start_,*t*_start_ + 0.25] s and converted to firing rates. Responses were averaged by stimulus position (*x, y*) to generate a 2D RF matrix, which was subsequently smoothed using a 2D Gaussian filter (*σ* = 1). To assess the spatial significance of the RF, we performed a permutation test (1,000 iterations). For each permutation, response values were spatially shuffled, and a *χ*^2^ statistic was computed to measure the deviation of the observed mean position vector from the global mean. The *p*-value was derived from the empirical null distribution. The RF centroid was calculated as the weighted center of mass of the smoothed RF positive values, and the RF area was defined as the pixel count of the suprathreshold mask (threshold set to 50% of the peak response).

#### Visual responsiveness and orientation selectivity

Visual responsiveness was quantified using peri-stimulus time histograms (PSTH, 10 ms bins) aligned to flash or grating onsets. Baseline and stimulus windows were defined as [− 0.25, 0] s and [0, 0.1] s, respectively. A unit was classified as visually responsive if the stimulus-evoked firing rate differed significantly from the baseline rate, assessed via paired Wilcoxon signed-rank test.

For orientation selectivity analysis, drifting grating responses were averaged across trials for each temporal frequency and orientation. The temporal frequency eliciting the maximum global Orientation Selectivity Index (gOSI) was selected as the optimal frequency. The gOSI was calculated using the vector sum of responses:

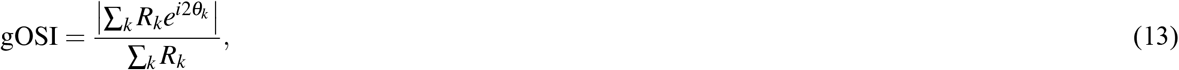

where *R*_*k*_ represents the rectified mean firing rate (stimulus rate minus baseline rate, negative values clamped to zero) at orientation *θ*_*k*_. The preferred orientation (PO) was derived from the phase of the resultant vector.

#### Ocular dominance analysis

We analysed ocular dominance (OD) in V1 units identified via channel mapping. Trials were aligned to stimulus onset with a baseline window of [− 1, 0] s and a stimulus window of [0, 1.5] s. We first determined the “best direction” for each eye independently. The best direction was defined as the direction with the highest stimulus firing rate that satisfied two criteria: (1) the stimulus rate exceeded the baseline rate, and (2) the difference was statistically significant (Mann-Whitney U test, *p <* 0.05). We calculated the baseline-subtracted evoked responses at the best direction for the contralateral (*C*) and ipsilateral (*I*) eyes. The Ocular Dominance Index (ODI) was defined as:

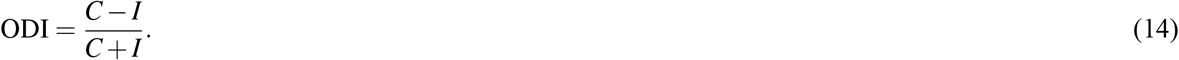

Based on ODI values, units were assigned to nine discrete categories using the boundaries [−1, −0.7, −0.5, −0.3, −0.1, 0.1, 0.3, 0.5, 0.7, 1]. Neurons in the first category are strongly dominated by ipsilateral responses, whereas those in the ninth category are strongly dominated by contralateral responses.

#### Binocular integration

Binocular integration was examined by comparing responses under binocular stimulation 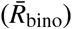 with monocular responses 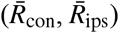. Unlike the OD analysis, response magnitudes for integration analysis were calculated by averaging the firing rates across *all* stimulus directions to capture the global integration property. The linear summation prediction was defined as 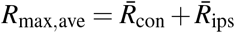. The Binocular Integration Ratio (BIR) was computed as:

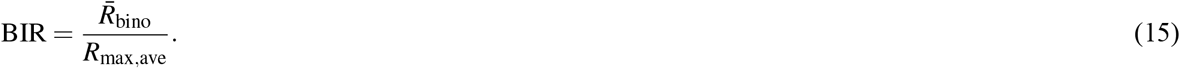

To avoid outliers, units with excessive firing rates (*>* 50 Hz) were excluded. Units were binned by their summed monocular response (*R*_max,ave_), and a one-sample *t*-test was performed within each bin to test if the BIR differed significantly from linearity (BIR = 1).

#### Human LFP event detection

LFP signals were averaged across all channels to obtain a global trace.

##### Burst-Suppression

We employed a recursive estimator for the running mean *µ*_*t*_ and variance 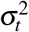 with a time constant *τ* = 0.5 s as described in Ref.^69^:

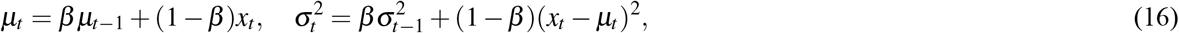

where β = exp[−1*/*(*f*_*s*_*τ*)]. Periods where 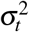 fell below a silence threshold were labelled as suppression. Burst onsets were defined as transitions from suppression to high-variance states, with a minimum burst duration of 0.5 s.

##### Interictal Epileptiform Discharges (IIDs)

IIDs were detected based on the algorithm described in Ref.^70^. The LFP was band-pass filtered (10-60 Hz), and the Hilbert envelope was extracted. An adaptive threshold was computed using a sliding window (5 s) as *k* × (mode + median) of the envelope distribution (*k* = 6.5). IIDs were identified as peaks exceeding this local threshold, with events merged if they occurred within 120 ms.

#### Spike-LFP coupling

##### IID Coupling

IID-locking was assessed by comparing firing rates in the event window [−0.1, +0.2] s against a baseline derived from non-event intervals. An ROC analysis was performed to quantify the discriminability of IID presence based on single-unit spike counts, reporting the Area Under the Curve (AUC).

##### Burst Coupling

A Burst Suppression Index (BSI) for single neurons was calculated using the firing rates during the burst phase (*R*_burst_, [0, 1] s) and the subsequent suppression phase (*R*_supp_, [1, 2] s) as 1 − *R*_supp_*/*(*R*_burst_).

#### Population Trajectory analysis

##### Human Neuropixels recordings

Spike trains were binned at 50 ms, trial-averaged, and z-scored to obtain normalized population activity matrices (time × neurons). To visualize low-dimensional dynamics, the population activity was smoothed with a Gaussian kernel (*σ* = 50 ms) and upsampled via cubic interpolation to 5 ms resolution. For IID events, Principal Component Analysis (PCA) was performed separately on responsive and non-responsive subpopulations to compare their state-space trajectories within a [−2.5, 2.5] s window relative to event onset. For burst events, PCA was applied to the population activity within a [−0.5, 1.5] s window.

##### IBL decision-making task

Neural trajectories were constructed from GRN population activity aligned to movement onset (*t* = 0). Instantaneous firing rates were computed using a sliding window (rectangular kernel; 12.5 ms width, 2 ms step) over the pre-movement interval [−0.15, 0] s. Firing rates were z-scored within each unit across time bins and baseline-subtracted (using [− 0.15, −0.14] s as baseline). To control for potential behavioral and task confounds, we balanced trial counts between left and right choices within strata defined by block prior and stimulus side via random subsampling. Choice-conditioned mean trajectories were computed by averaging across balanced trials. PCA was performed on the concatenated time-by-neuron matrix (stacking left and right trajectories; samples = 2*N*_*t*_, features = *N*_unit_). The first three PCs were used to visualize 3D trajectories, with the sign of PC1 flipped to enforce a uniform temporal direction.

#### Decoding analysis

##### IBL behavioral Decoding

For the IBL decision-making task, we decoded the animal’s choice from population activity in the GRN. Firing rates in the pre-movement window [−0.1, 0] s were used as features. A Logistic Regression classifier with L1 regularization was trained using a nested cross-validation scheme (5-fold outer, 4-fold inner for hyperparameter search).

##### Human IID Decoding

A Random Forest classifier (200 trees) was trained to distinguish IID event windows from baseline windows based on population PSTH features. Performance was evaluated using AUROC computed over 50 bootstrap iterations.

#### Statistical analysis

Statistical analyses were performed using Python, primarily with the SciPy and scikit-learn packages. Unless otherwise stated, quantitative data are presented as mean ± standard error of the mean (SEM). Data normality was assessed using the Shapiro-Wilk test. For normally distributed data, comparisons between two groups were performed using Student’s *t*-test (paired or unpaired as appropriate), and comparisons against a fixed reference value were performed using one-sample *t*-tests. For non-normally distributed data, non-parametric tests were used, including the Mann-Whitney U test for unpaired comparisons and the Wilcoxon signed-rank test for paired comparisons. Categorical data distributions were compared using Fisher’s exact test, and correlations were assessed using Spear-man’s rank correlation coefficient. When multiple comparisons were performed within a single analysis, *p*-values were adjusted using either Bonferroni correction or Benjamini-Hochberg false discovery rate control, as specified in the corresponding figure legends. Statistical significance was defined as *p <* 0.05.

## Resource availability

## Data availability

This study used several publicly available datasets, including the Hybrid Neuropixels dataset (https://janelia.figshare.com/articles/dataset/Simulations_from_kilosort4_paper/25298815/1), the Allen Brain Observatory Visual Coding dataset (https://allensdk.readthedocs.io/en/latest/visual_coding_neuropixels.html), the International Brain Laboratory (IBL) decision-making dataset (https://viz.internationalbrainlab.org), and the Human Neuropixels dataset (https://datadryad.org/dataset/doi:10.5061/dryad.d2547d840).

Source data required to reproduce the figures are provided with this manuscript. All other data supporting the findings of this study are available from the corresponding author upon reasonable request.

## Code availability

All custom code used in this study, including framework pipelines, training and inference code, pretrained model weights, and the analysis scripts for each figure, is provided with this manuscript.

## Acknowledgements

This study was supported by Microsoft Research and the National Natural Science Foundation of China (32471055 and 82171090 to Y.G.), AI for Science Foundation of Fudan University (FudanX24AI046 to Y.G.), ZJLab, Shanghai Center for Brain Science and Brain-Inspired Technology, the Lingang Laboratory.

## Author contributions

Y.Z. and D.H. conceived the study, designed and implemented the computational framework, and performed model training and evaluation. Y.Z. performed electrophysiology simulations. Y.Z. and Z.L. performed data and statistical analysis. Y.G. contributed to the interpretation of the results. Z.L. and F.R. contributed to data acquisition, curation, and analysis. Y.Z., D.H., and Y.G. wrote the manuscript with input from all authors. D.L., Y.Y., and Y.W. provided experimental resources and contributed to project discussions. Y.G. and D.L. supervised the project and acquired funding. All authors read and approved the final manuscript.

## Declaration of interests

The authors declare no competing interests.

---

**Figure S1.**
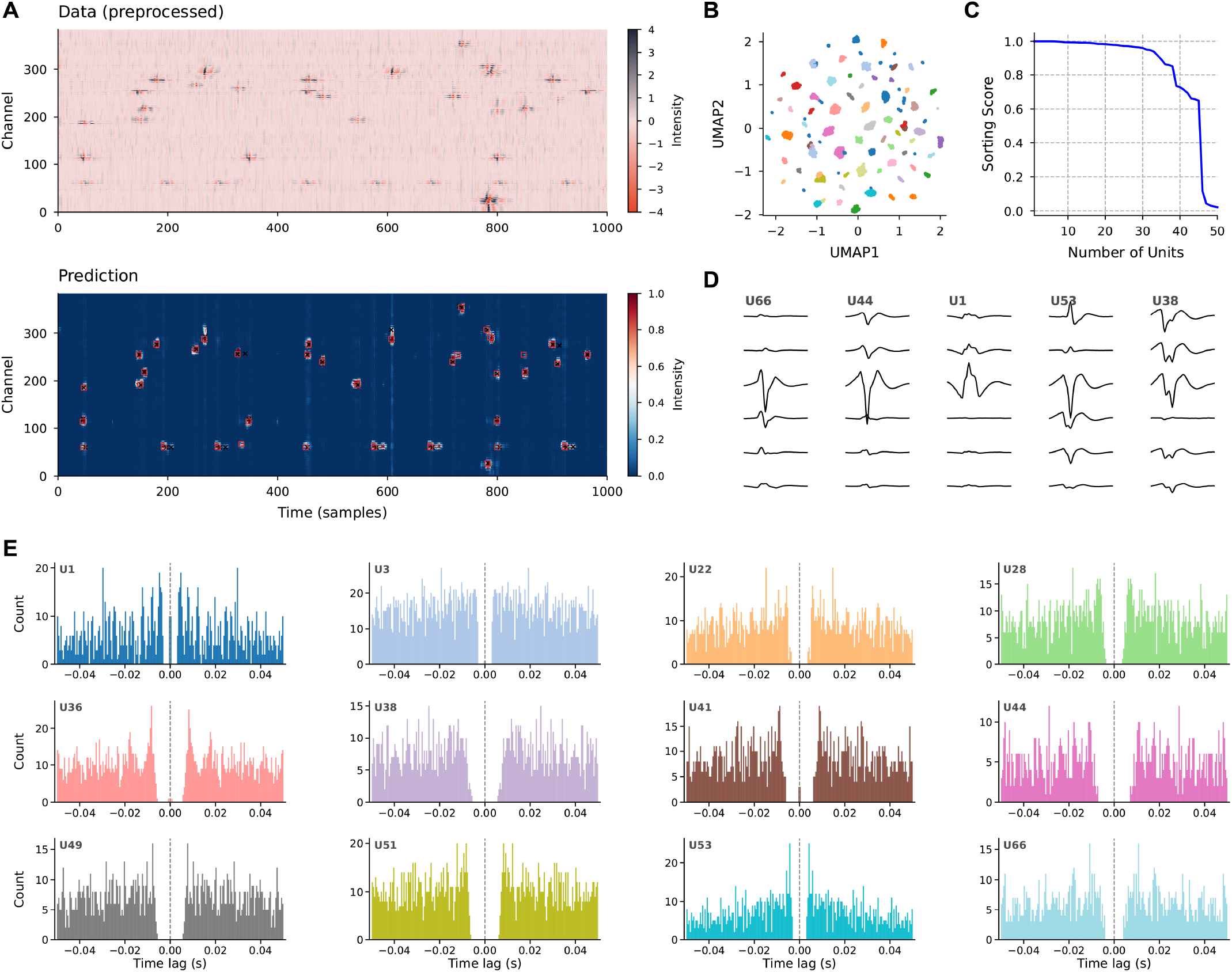
Visualization of model-inferred sorting results on BBP-L6 dataset. **(A)** Visualization of detection inference on the simulated recording. The top panel shows the preprocessed data, and the bottom panel shows the model’s predicted probability map, where red squares indicate ground truth spikes and black crosses indicate detected peaks. **(B)** Visualization of spike waveform features projected into a 2D UMAP embedding space, coloured by their assigned cluster identities. **(C)** Sorting performance distribution curve, plotting the sorting F1 scores for all sorted units ranked in descending order. **(D)** Spatially distributed waveform footprints of representative sorted units across main channels. **(E)** Autocorrelograms of example units, demonstrating clear refractory periods indicative of single-unit isolation.

**Figure S2.**
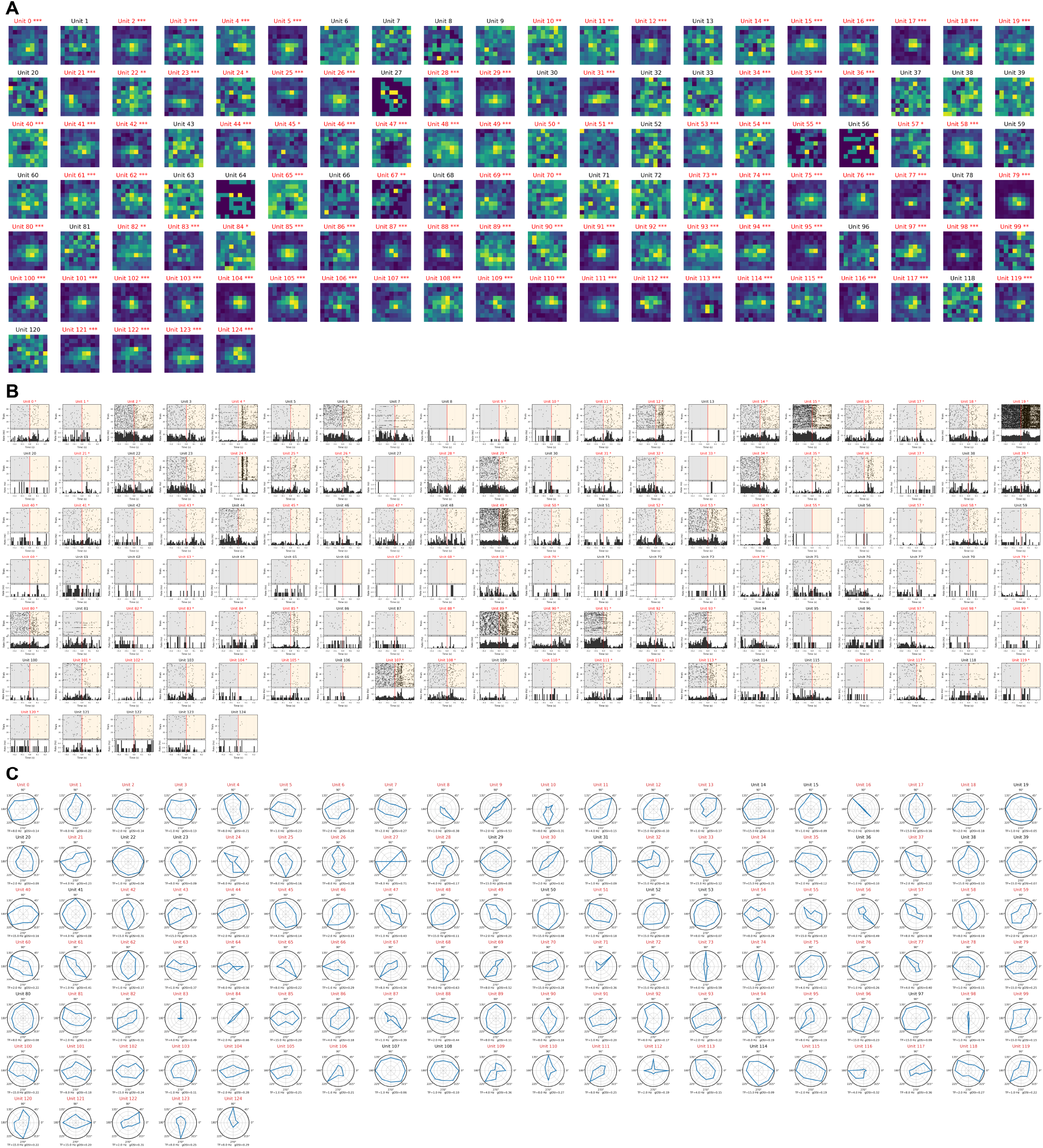
Visual response properties of model-inferred units from the Allen Visual Coding dataset. **(A)** Spatial receptive fields mapped using Gabor stimuli. Heatmaps illustrate the mean firing rate at each stimulus position (*x, y*), smoothed with a 2D Gaussian filter. Asterisks denote significant spatial structure (permutation test). **(B)** Visual responses to full-field flash stimuli. Top panels: Raster plots aligned to stimulus onset (red line), with OFF (grey) and ON (beige) periods indicated. Bottom panels: PSTHs showing mean firing rate. Asterisks indicate significant deviations from baseline (p < 0.05, bootstrap test). **(C)** Orientation tuning properties. Polar plots display mean firing rates as a function of drifting grating direction at the optimal temporal frequency. Unit titles coloured red indicate *gOSI >* 0.1.

**Figure S3.**
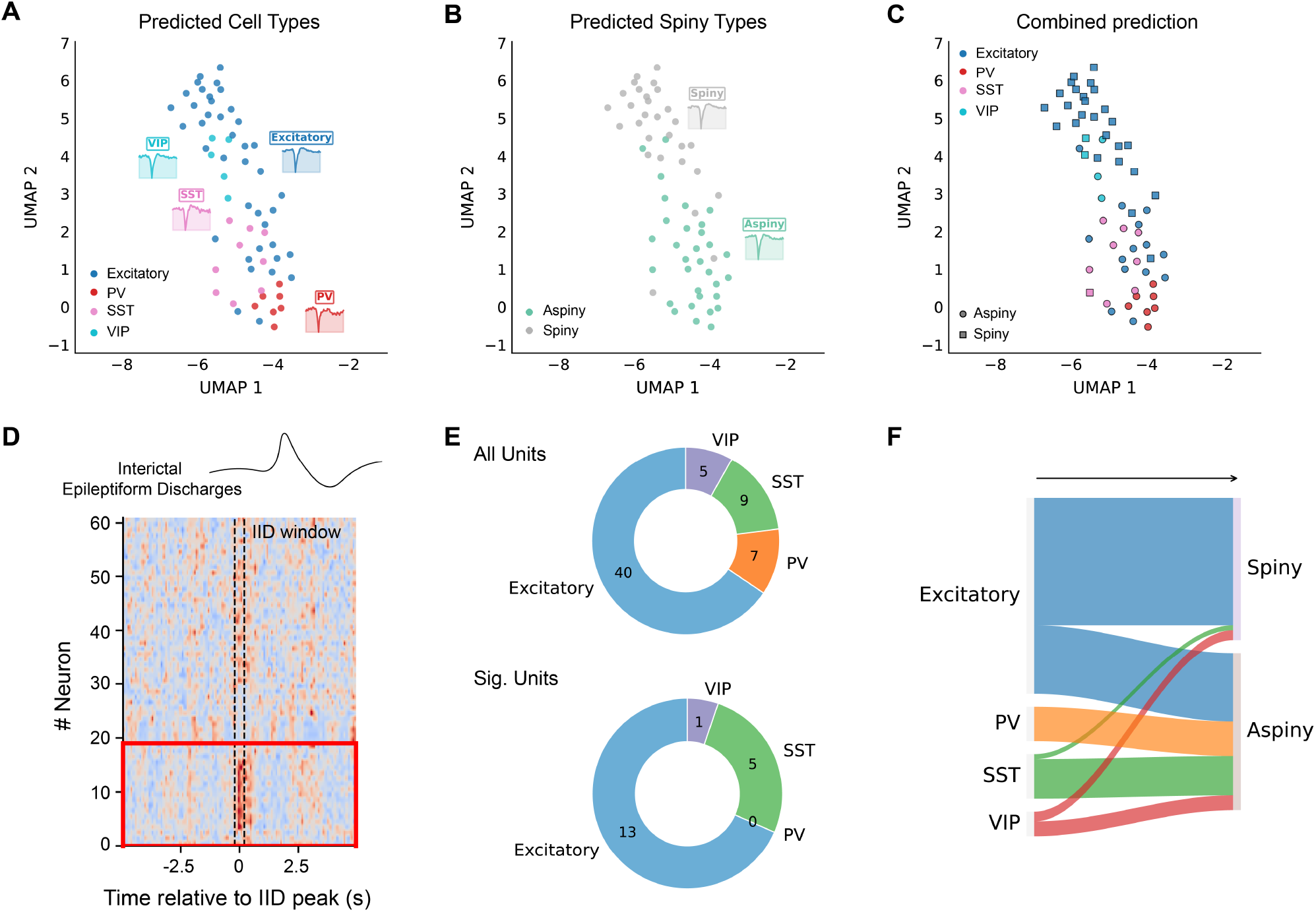
Prediction of cell identities and spiny morphologies in human cortical recordings. **(A-C)** UMAP visualization of the latent feature space learned by the cell classification model. Each dot represents a single unit. The embedding space is coloured by predicted cell types in **A** (insets: representative average waveforms) and by predicted spiny types in **B. (C)** Joint visualization of predicted cell types (color) and spiny morphologies (shape; square: Spiny, circle: Aspiny), showing the correspondence between the two classifications. **(D)** Heatmap of population firing rates aligned to the peak of interictal epileptiform discharges (IIDs), corresponding to Figure 5I. **(E)** Distribution of predicted cell types within the total recorded population (top) versus the sub-population significantly modulated by IIDs (bottom). **(F)** Sankey diagram illustrating the mapping concordance between predicted cell identities and spiny morphologies.

**Figure S4.**
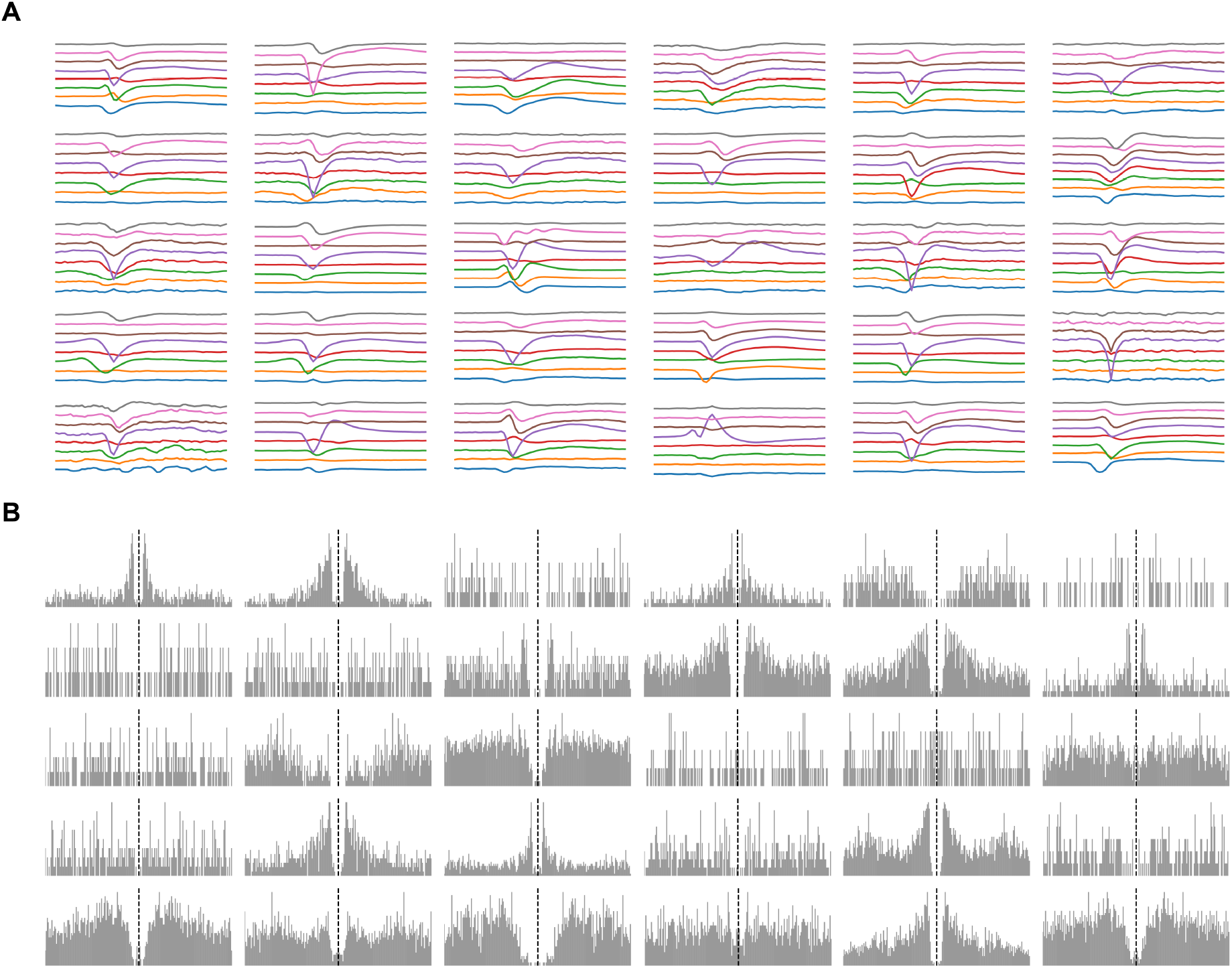
Example weak-response neurons in Figure 6. **(A)** Extracellular waveforms of representative weak-response units spanning the main recording channels. **(B)** Corresponding autocorrelograms revealing distinct refractory periods characteristic of single-unit isolation.

## Notes

### Competing Interest Statement

The authors have declared no competing interest.

## References

1. Rall, W. Electrophysiology of a Dendritic Neuron Model. Biophys. J. 2, 145–167 (1962).

2. Hodgkin, A. L. & Huxley, A. F. A quantitative description of membrane current and its application to conduction and excitation in nerve. Bull. Math. Biol. 52, 25–71 (1990).

3. Destexhe, A., Mainen, Z. F. & Sejnowski, T. J. Synthesis of models for excitable membranes, synaptic transmission and neuromodulation using a common kinetic formalism. J. Comput. Neurosci. 1, 195–230 (1994).

4. Gerstner, W. & Kistler, W. M. Spiking Neuron Models: Single Neurons, Populations, Plasticity (Cambridge University Press, 2002).

5. Markram, H. et al. Reconstruction and Simulation of Neocortical Microcircuitry. Cell 163, 456–492 (2015).

6. Billeh, Y. N. et al. Systematic Integration of Structural and Functional Data into Multi-scale Models of Mouse Primary Visual Cortex. Neuron 106, 388–403.e18 (2020).

7. Steinmetz, N. A. et al. Neuropixels 2.0: A miniaturized high-density probe for stable, long-term brain recordings. Science 372, eabf4588 (2021).

8. Ye, Z. et al. Ultra-high-density neuropixels probes improve detection and identification in neuronal recordings. Neuron 113, 3966–3982.e12 (2025).

9. Stringer, C. et al. Spontaneous behaviors drive multidimensional, brainwide activity. Science 364, eaav7893 (2019).

10. Siegle, J. H. et al. Survey of spiking in the mouse visual system reveals functional hierarchy. Nature 592, 86–92 (2021).

11. Pachitariu, M., Steinmetz, N. A., Kadir, S. N., Carandini, M. & Harris, K. D. Fast and accurate spike sorting of high-channel count probes with KiloSort. In Advances in Neural Information Processing Systems, vol. 29 (Curran Associates, Inc., 2016).

12. Chung, J. E. et al. A Fully Automated Approach to Spike Sorting. Neuron 95, 1381–1394.e6 (2017).

13. Hilgen, G. et al. Unsupervised Spike Sorting for Large-Scale, High-Density Multielectrode Arrays. Cell Reports 18, 2521–2532 (2017).

14. Buccino, A. P., Garcia, S. & Yger, P. Spike sorting: New trends and challenges of the era of high-density probes. Prog. Biomed. Eng. 4, 022005 (2022).

15. Pachitariu, M., Sridhar, S., Pennington, J. & Stringer, C. Spike sorting with Kilosort4. Nat. Methods 21, 914–921 (2024).

16. Laquitaine, S., Imbeni, M., Tharayil, J., Isbister, J. B. & Reimann, M. W. Spike sorting biases and information loss in a detailed cortical model. bioRxiv 2024.12.04.626805 (2025).

17. Lee, E. K. et al. PhysMAP - interpretable in vivo neuronal cell type identification using multi-modal analysis of electrophysiological data. Neuroscience (2024).

18. Lee, E. K. et al. Non-linear dimensionality reduction on extracellular waveforms reveals cell type diversity in premotor cortex. eLife 10, e67490 (2021).

19. Trainito, C., von Nicolai, C., Miller, E. K. & Siegel, M. Extracellular Spike Waveform Dissociates Four Functionally Distinct Cell Classes in Primate Cortex. Curr. Biol. 29, 2973–2982.e5 (2019).

20. Wei, Y. et al. Associations between in vitro, in vivo and in silico cell classes in mouse primary visual cortex. Nat. Commun. 14, 2344 (2023).

21. Degrave, J. et al. Magnetic control of tokamak plasmas through deep reinforcement learning. Nature 602, 414–419 (2022).

22. Bellemare, M. G. et al. Autonomous navigation of stratospheric balloons using reinforcement learning. Nature 588, 77–82 (2020).

23. Kaufmann, E. et al. Champion-level drone racing using deep reinforcement learning. Nature 620, 982–987 (2023).

24. Quian Quiroga, R. & Panzeri, S. Extracting information from neuronal populations: Information theory and decoding approaches. Nat. Rev. Neurosci. 10, 173–185 (2009).

25. Harris, K. D., Henze, D. A., Csicsvari, J., Hirase, H. & Buzsáki, G. Accuracy of Tetrode Spike Separation as Determined by Simultaneous Intracellular and Extracellular Measurements. J. Neurophysiol. (2000).

26. Barnett, A. H., Magland, J. F. & Greengard, L. F. Validation of neural spike sorting algorithms without ground-truth information. J. Neurosci. Methods 264, 65–77 (2016).

27. Einevoll, G. T., Kayser, C., Logothetis, N. K. & Panzeri, S. Modelling and analysis of local field potentials for studying the function of cortical circuits. Nat. Rev. Neurosci. 14, 770–785 (2013).

28. Hay, E., Hill, S., Schürmann, F., Markram, H. & Segev, I. Models of Neocortical Layer 5b Pyramidal Cells Capturing a Wide Range of Dendritic and Perisomatic Active Properties. PLOS Comput. Biol. 7, e1002107 (2011).

29. Gouwens, N. W. et al. Systematic generation of biophysically detailed models for diverse cortical neuron types. Nat. Commun. 9, 710 (2018).

30. Hines, M. L. & Carnevale, N. T. The NEURON Simulation Environment. Neural Comput. 9, 1179–1209 (1997).

31. Hines, M. L. & Carnevale, N. T. Neuron: A Tool for Neuroscientists. The Neurosci. 7, 123–135 (2001).

32. Lindén, H. et al. LFPy: A tool for biophysical simulation of extracellular potentials generated by detailed model neurons. Front. Neuroinformatics 7 (2014).

33. Jun, J. J. et al. Fully integrated silicon probes for high-density recording of neural activity. Nature 551, 232–236 (2017).

34. International Brain Laboratory et al. Reproducibility of in vivo electrophysiological measurements in mice. eLife 13, RP100840 (2025).

35. de Vries, S. E., Siegle, J. H. & Koch, C. Sharing neurophysiology data from the Allen Brain Observatory. eLife 12, e85550 (2023).

36. Steinmetz, N. A., Zatka-Haas, P., Carandini, M. & Harris, K. D. Distributed coding of choice, action and engagement across the mouse brain. Nature 576, 266–273 (2019).

37. Meshulam, L. et al. A brain-wide map of neural activity during complex behaviour. Nature 645, 177–191 (2025).

38. Hubel, D. H. & Wiesel, T. N. Receptive fields, binocular interaction and functional architecture in the cat’s visual cortex. The J. Physiol. 160, 106–154 (1962).

39. Niell, C. M. & Stryker, M. P. Highly Selective Receptive Fields in Mouse Visual Cortex. J. Neurosci. 28, 7520–7536 (2008).

40. Allen Institute for Brain Science. Allen cell types database. https://celltypes.brain-map.org (2015).

41. Hodge, R. D. et al. Conserved cell types with divergent features in human versus mouse cortex. Nature 573, 61–68 (2019).

42. Paulk, A. C. et al. Large-scale neural recordings with single neuron resolution using Neuropixels probes in human cortex. Nat. Neurosci. 25, 252–263 (2022).

43. Cang, J., Fu, J. & Tanabe, S. Neural circuits for binocular vision: Ocular dominance, interocular matching, and disparity selectivity. Front. Neural Circuits 17, 1084027 (2023).

44. Gordon, J. A. & Stryker, M. P. Experience-Dependent Plasticity of Binocular Responses in the Primary Visual Cortex of the Mouse. J. Neurosci. 16, 3274–3286 (1996).

45. Fagiolini, M. & Hensch, T. K. Inhibitory threshold for critical-period activation in primary visual cortex. Nature 404, 183–186 (2000).

46. McGee, A. W., Yang, Y., Fischer, Q. S., Daw, N. W. & Strittmatter, S. M. Experience-driven plasticity of visual cortex limited by myelin and nogo receptor. Science 309, 2222–2226 (2005).

47. Morishita, H., Miwa, J. M., Heintz, N. & Hensch, T. K. Lynx1, a cholinergic brake, limits plasticity in adult visual cortex. Science 330, 1238–1240 (2010).

48. Salinas, K. J., Velez, D. X. F., Zeitoun, J. H., Kim, H. & Gandhi, S. P. Contralateral Bias of High Spatial Frequency Tuning and Cardinal Direction Selectivity in Mouse Visual Cortex. J. Neurosci. 37, 10125–10138 (2017).

49. Huh, C. Y. L. et al. Long-term monocular deprivation during juvenile critical period disrupts binocular integration in mouse visual thalamus. J. Neurosci. 40, 585–604 (2020).

50. Jenks, K. R. & Shepherd, J. D. Experience-dependent development and maintenance of binocular neurons in the mouse visual cortex. Cell Reports 30, 1982–1994.e4 (2020).

51. Tan, L., Tring, E., Ringach, D. L., Zipursky, S. L. & Trachtenberg, J. T. Vision changes the cellular composition of binocular circuitry during the critical period. Neuron 108, 735–747.e6 (2020).

52. Zhao, X., Liu, M. & Cang, J. Sublinear binocular integration preserves orientation selectivity in mouse visual cortex. Nat. Commun. 4, 2088 (2013).

53. Garrett, M. E., Nauhaus, I., Marshel, J. H. & Callaway, E. M. Topography and Areal Organization of Mouse Visual Cortex. The J. Neurosci. 34, 12587–12600 (2014).

54. Quiroga, R., Nadasdy, Z. & Ben-Shaul, Y. Unsupervised spike detection and sorting with wavelets and super-paramagnetic clustering. Neural Comput. 16, 1661–1687 (2004).

55. Rossant, C. et al. Spike sorting for large, dense electrode arrays. Nat. Neurosci. 19, 634–641 (2016).

56. Tobin, J. et al. Domain randomization for transferring deep neural networks from simulation to the real world. In 2017 IEEE/RSJ International Conference on Intelligent Robots and Systems (IROS), 23–30 (2017).

57. Peng, X. B., Andrychowicz, M., Zaremba, W. & Abbeel, P. Sim-to-Real Transfer of Robotic Control with Dynamics Randomization. In 2018 IEEE International Conference on Robotics and Automation (ICRA), 3803–3810 (2018).

58. Ramaswamy, S. et al. The neocortical microcircuit collaboration portal: A resource for rat somatosensory cortex. Front. Neural Circuits 9 (2015).

59. Hagen, E. et al. ViSAPy: A Python tool for biophysics-based generation of virtual spiking activity for evaluation of spike-sorting algorithms. J. Neurosci. Methods 245, 182–204 (2015).

60. Hagen, E., Næss, S., Ness, T. V. & Einevoll, G. T. Multimodal Modeling of Neural Network Activity: Computing LFP, ECoG, EEG, and MEG Signals With LFPy 2.0. Front. Neuroinformatics 12 (2018).

61. Buccino, A. P. & Einevoll, G. T. MEArec: A Fast and Customizable Testbench Simulator for Ground-truth Extracellular Spiking Activity. Neuroinformatics 19, 185–204 (2021).

62. Holmes, W. R. Passive Cable Modeling. In Computational Modeling Methods for Neuroscientists (2009).

63. Brette, R. & Destexhe, A. Handbook of Neural Activity Measurement. In Handbook of Neural Activity Measurement, 92–135 (Cambridge University Press, 2012).

64. Buzsáki, G., Anastassiou, C. A. & Koch, C. The origin of extracellular fields and currents —EEG, ECoG, LFP and spikes. Nat. Rev. Neurosci. 13, 407–420 (2012).

65. Schroff, F., Kalenichenko, D. & Philbin, J. FaceNet: A unified embedding for face recognition and clustering. In 2015 IEEE Conference on Computer Vision and Pattern Recognition (CVPR), 815–823 (2015).

66. Zhang, Y. et al. SimSort: A Data-Driven Framework for Spike Sorting by Large-Scale Electrophysiology Simulation. In The Thirty-ninth Annual Conference on Neural Information Processing Systems (2025).

67. Loshchilov, I. & Hutter, F. Decoupled weight decay regularization. In International Conference on Learning Representations (2018).

68. Kingma, D. P. & Ba, J. Adam: A Method for Stochastic Optimization. In International Conference on Learning Representations (2014).

69. Brandon Westover, M. et al. Real-time segmentation of burst suppression patterns in critical care EEG monitoring. J. Neurosci. Methods 219, 131–141 (2013).

70. Janca, R. et al. Detection of interictal epileptiform discharges using signal envelope distribution modelling: Application to epileptic and non-epileptic intracranial recordings. Brain Topogr. 28, 172–183 (2015).

